# Donut-like organization of inhibition underlies categorical neural responses in the midbrain

**DOI:** 10.1101/2019.12.24.887810

**Authors:** Nagaraj R. Mahajan, Shreesh P. Mysore

## Abstract

Categorical neural responses underlie various forms of selection and decision-making. Such binary-like responses promote robust signaling of the winner in the presence of input ambiguity and neural noise. Here, we show that a ‘donut-like’ inhibitory mechanism in which each competing option suppresses all options except itself, is highly effective at generating categorical neural responses. It surpasses motifs of feedback inhibition, recurrent excitation, and divisive normalization invoked frequently in decision-making models. We demonstrate experimentally not only that this mechanism operates in the midbrain spatial selection network in barn owls, but also that it is necessary for categorical signaling by it. The functional pattern of neural inhibition in the midbrain forms an exquisitely structured ‘multi-holed’ donut consistent with this network’s combinatorial inhibitory function for stimulus selection. Additionally, modeling reveals a generalizable neural implementation of the donut-like motif for categorical selection. Self-sparing inhibition may, therefore, be a powerful circuit module central to categorization.

## INTRODUCTION

Categorization, the transformation of continuously varying inputs into discrete output groups, is a fundamental component of perception and decision-making^1–3^. Neural responses that are explicitly categorical^4^, have been reported across brain areas and animal species in a variety of perceptual and decision-making contexts ^2,5–13^. Such response profiles, which involve a large and abrupt change in firing rate across the category boundary (Fig. 1A-left, red), are computationally advantageous: they enhance downstream decoding of the selected category (or ‘winner’) particularly when competing options are similar (i.e., in the face of input ambiguity), and when neural responses are variable (i.e., in the face of representational uncertainty; Fig. 1A; red vs. blue) ^9,14^. Despite the pervasiveness ^2,5–13^ and computational benefit of such categorical neural response profiles, how the brain implements them is unknown. Specifically, it is unclear what identifiable circuit mechanisms are essential for producing categorical neural response profiles.

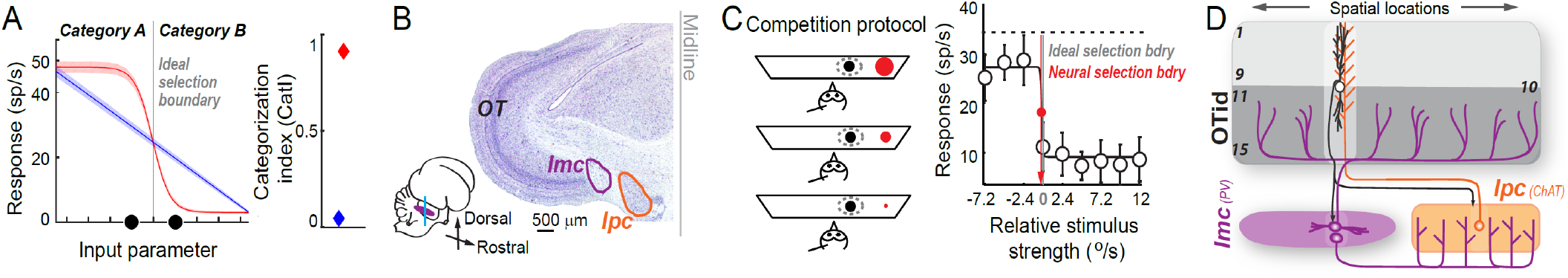
**(B)** Midbrain selection network in barn owl. <u>Inset</u>: Side view of owl brain. Vertical blue line: line of section. <u>Main</u>: Coronal section through owl midbrain showing optic tectum (OT), isthmi pars magnocellularis (Imc, GABAergic, parvalbumin-positive, purple outline), and isthmi pars parvocellularis (Ipc, cholinergic, orange outline). **(C)** <u>Left:</u> Schematic of “competition protocol” used in (non-behaving) owls^9,22^. A stimulus of fixed strength (black dot) is presented inside the spatial receptive field (RF; dashed oval) of a neuron, while a competitor (red dot) of varying strength is presented far outside the RF. Strength of the stimuli is controlled by their loom speeds (°/s); denoted here by size of dots. <u>Right:</u> Categorical (‘switch-like’) response profile of a neuron in the intermediate and deep layers of the OT (OTid) measured using the competition protocol in left panel. Red vertical line: neural selection boundary; indicates the relative strength at which neural responses switch from being at a high level to a low level. Gray vertical line: ideal selection boundary; indicates the relative strength of 0, at which the two stimuli are equally strong. The neural selection boundary nearly overlaps with ideal one. Data reproduced with permission^9^. **(D)** Schematic of connectivity within the avian midbrain selection network. Layers (1-15) of OT are shown. OTid: intermediate and deep layers (layers 11-15). Cholinergic Ipc neuron (orange circle) receives focal input from OT (black circle, layer 10), and send projections focally back to OT (orange projections). GABAergic Imc neuron (purple circles) receives focal input from OT (black circle, layer 10), and but sends inhibitory projections broadly across OTid as well as Ipc (purple projections). Consequently, Imc neurons suppress OTid responses via two pathways: (i) the direct pathway to the OTid^31^ (Imc→OT), and (ii) the indirect pathway that inhibits the potent point-to-point cholinergic amplifiers of OTid, namely the Ipc^31,33^ (Imc→ Ipc→ OT). **See also Fig. S1**.

An excellent site in the brain at which to investigate neural circuit mechanisms of categorization is a vertebrate midbrain network that plays a causal role in controlling gaze and spatial attention ^15–19 20,21^. This network, which encodes sensory space topographically, includes the optic tectum (called the superior colliculus, SC, in mammals) and several nuclei in the midbrain tegmentum, referred to as the isthmic nuclei ^21^(Fig. 1B). In the barn owl, this network has been shown to categorize stimuli into two categories: ‘‘highest priority” and ‘‘others’’^8,9,22^, with the priority of a stimulus being defined as the combination of its physical salience and behavioral relevance ^23^. This categorization manifests as ‘switch-like’ responses in a subset of neurons in the intermediate and deep layers of the owl optic tectum (OTid, SCid in mammals) ^8,9,22^. These neurons fire at a high rate when the stimulus inside their spatial receptive field (RF) is the highest priority, but switch abruptly to a lower firing rate when a distant, competing stimulus becomes the highest priority one (Fig. 1C; red neural selection boundary nearly overlaps with grey ideal selection boundary). Such switch-like responses markedly improve discriminability of the location of the highest priority stimulus among competing stimuli of similar priority ^9,14^. Additionally, such categorization by OTid in owls accounts well for the specific pattern of spatial selection deficits observed in monkeys following inactivation of the SCid^14^: worsening of the impairment in selecting a target among distracters as they become more similar to the target ^17,18,24^. Together, these studies support that categorical responses in OTid enhance reliable readout of the highest priority stimulus for gaze or spatial attention behavior, particularly when competing stimuli are of similar strength (or more generally, similar priority)^14,21,22^.

The responses of OTid neurons are regulated by two key isthmic nuclei in the midbrain selection network. OTid responses to single stimuli are enhanced multiplicatively by cholinergic neurons of the isthmi pars parvocellularis (Ipc), which exhibit point-to-point recurrent connectivity with the OT (Fig. 1D-orange)^25–27^. In parallel, responses of OTid neurons to multiple competing stimuli are controlled by inhibitory neurons of the isthmi pars magnocellularis (Imc) (Fig. 1D-purple) ^28–31^. Imc neurons receive focal input from the OT but send long-range projections broadly across both the OTid and Ipc space maps, suppressing OT through the direct (Imc→OTid) pathway as well as the indirect (Imc→Ipc→OTid) pathway (Fig. 1D-purple projections). The Imc controls competitive interactions across the OTid space map: focal inactivation of Imc neurons abolishes all competitive interactions in the OTid (and Ipc) ^32,33^. Despite these insights, how categorical neural response profiles in the OTid are generated is an open question.

Here, using a combination of neurally grounded computational modeling, dual electrophysiological recordings, and focal iontophoretic neural inactivation, we investigate how categorical neural responses in the barn owl OTid are generated by computations in the OT-Imc-Ipc network. We demonstrate that a donut-like pattern of spatial inhibition in this network causally controls categorization by OTid, and also that this inhibition is implemented with an intricate multi-holed donut-like pattern across interconnected brain areas in support of categorization across space. These results suggest that donut-like inhibition may be a fundamental circuit motif for generating categorical neural responses, a function central to spatial selection, perception, and decision making ^1,34^.

## RESULTS

### Donut-like inhibitory motif emerges as a powerful mechanism for generating categorical responses in model circuits of the avian midbrain selection network

As a first step in investigating the circuit mechanisms underlying categorical neural response profiles in the OTid, we turned to neurally grounded computational modeling. Starting with a simple model of the midbrain network capable of comparing the representations of competing stimuli^35–37^ (‘Baseline’ model; Fig. 2A, left column-top, oval ‘Imc’ neurons deliver feedforward inhibition to circular ‘OTid’ neurons; Methods), we introduced, systematically, each of three circuit motifs that have been proposed in the literature as potential mechanisms for generating categorical response profiles^38–41^. We compared directly the ability of these different circuit models to generate categorical neural response profiles.

**Fig. 2.**
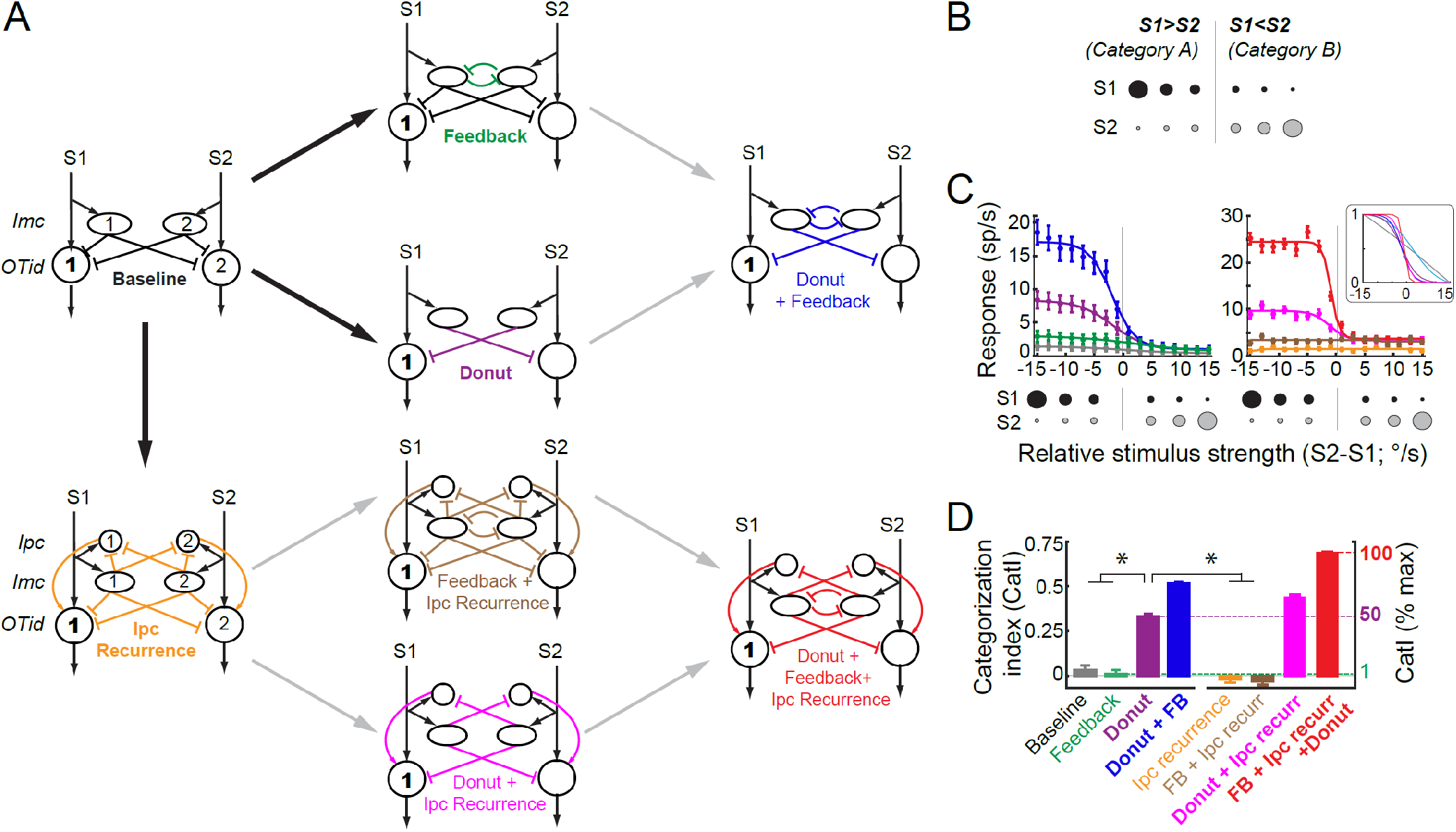
Donut-like inhibition surpasses other hypothesized circuit motifs for generating categorical responses in models of the avian midbrain selection network. **(A)** Computational models incorporating combinations of three different circuit motifs proposed in the literature as underlying categorical responses (Methods; Fig. S1). All models built upon generic ‘baseline’ circuit (left column-top) capable of comparing competing options. Each model shows two ‘channels’; each channel is the group of neurons (numbered) involved in representing a stimulus (S1 or S2). Large circles: OTid neurons; ovals: Imc neurons; small circles: Ipc neurons. Arrows with pointed heads - excitatory connections, arrows with flat heads - inhibitory connections. The three circuit motifs are – feedback inhibition between competing channels (green; middle column - top), donut-like inhibition (self-sparing inhibition; purple; middle column-second from top; see also text), and recurrent amplification within each channel (through ‘Ipc’ neuron; orange; left column-bottom). The goal of output neuron 1 (bold face) in each model is to signal if S1>S2 (category ‘a’) or S2>S1 (category ‘b’), when presented with S1 and S2 of varying relative strength following strength-morphing protocol. **(B)** Strength-morphing protocol (Methods). S1 and S2 are presented simultaneously ‘to’ the model, S1(2) is inside the receptive field of neurons in channel 1 (2). As strength of S1 is decreased, that of S2 is systematically increased; strength of the stimuli is controlled by their loom speeds (°/s); denoted here by size of dots. Gray line: ideal selection boundary (when relative strength =0) **(C)** Simulated response profiles of output neuron 1 from each of the models (colors) in A obtained using the strength-morphing protocol (bottom inset). Responses are mean ± s.e.m of 30 repetitions. The continuously varied input parameter was the relative strength of the two stimuli (S2-S1). Lines – best sigmoidal fits. Input-output functions of model neurons were sigmoids with Gaussian noise (Methods; fano factor=6). Right-Inset: Response profiles normalized between 0 and 1; only means are shown for clarity. **(D)** ‘Population’ summary of CatI of response profiles from various circuit models (colors); mean ± s.e.m; n=50 model neurons. ‘*’: p<0.05, ANOVA followed by Holm-Bonferroni correction for multiple paired comparisons; only a key subset of significant differences indicated for clarity. **See also Fig. S1.**

The first mechanistic proposal from published literature is that feedback inhibition between the representations of competing options plays a key role in categorization. This is motivated by modeling studies of categorical decision-making^40,42^, work on direction selectivity in the retina^43^, as well as work on spatial selection in barn owls^44^. The reasoning is that the iterative nature of the feedback inhibition may allow for small differences in competing options to be amplified, resulting in large differences in steady-state competitive inhibition, and therefore, in neural responses. In the avian midbrain network, such feedback inhibition is known to be implemented as long-range reciprocal inhibitory projections among Imc neurons representing different (competing) options ^31,44,45^. Therefore, in our model, we introduced feedback inhibition as reciprocal inhibition between the two model Imc neurons (Fig. 2A, middle-column top; green inhibitory connections)^44^.

The second mechanistic proposal is that recurrent excitation of the responses to each option plays a key role in categorization^40,46^. This is a common element in models of decision-making and is thought to aid categorical selection through response amplification ^40,46^. In the avian midbrain network, recurrent excitation of OTid responses encoding for a particular spatial location is known to be implemented by the cholinergic Ipc neurons encoding for the same location ^26^, resulting in focal multiplicative enhancement of OTid activity ^40,46^ (Fig. 1D). Notably, as mentioned previously, Ipc neurons receive feedforward inhibition from Imc (Fig. 1D)^26,31^, as a result of which recurrent amplification of OTid by Ipc is regulated by Imc. Therefore, in our model, we introduced ‘Ipc recurrence’ by incorporating amplifying Ipc neurons that lie downstream of Imc (Fig. 2A, left column-bottom; orange projections).

The third mechanistic proposal is that a donut-like pattern of competitive inhibition, i.e., one in which the representation of each option suppresses others more strongly than it suppresses itself, may play a key role in categorization. This is motivated by work in turtles^47^ as well as by the observation that categorical response profiles exhibit large response differences across the selection boundary. The reasoning is that having strong inhibition to ‘other’ options, but weak ‘self’-inhibition can enhance response differences. In the avian midbrain network, inhibitory projections from each Imc neuron, which span the OTid space maps broadly (Fig. 1D), are thought to spare the portion of OT from which the Imc neuron receives input, suggesting an anatomical donut-like pattern of inhibition in the direct pathway from Imc to OTid ^31^. Similar details about the indirect Imc→Ipc→ OT pathway are unknown. Therefore, in our model, we introduced donut-like inhibition by removing self-inhibitory connections in the direct Imc→OT pathway (Fig. 2A, middle column-second from top model; absence of purple projections from Imc neurons to aligned OTid neurons). (For completeness, in models that included Ipc recurrence, we also removed self-inhibitory connections in the indirect Imc→Ipc→OT pathway: Fig. 2A, middle column-bottom, absence of absence of pink projections from Imc neurons to aligned Ipc neurons; right column-bottom, absence of absence of red projections from Imc neurons to aligned Ipc neurons; see also Fig. 3 for direct experimental validation of this assumption.)

**Fig. 3.**
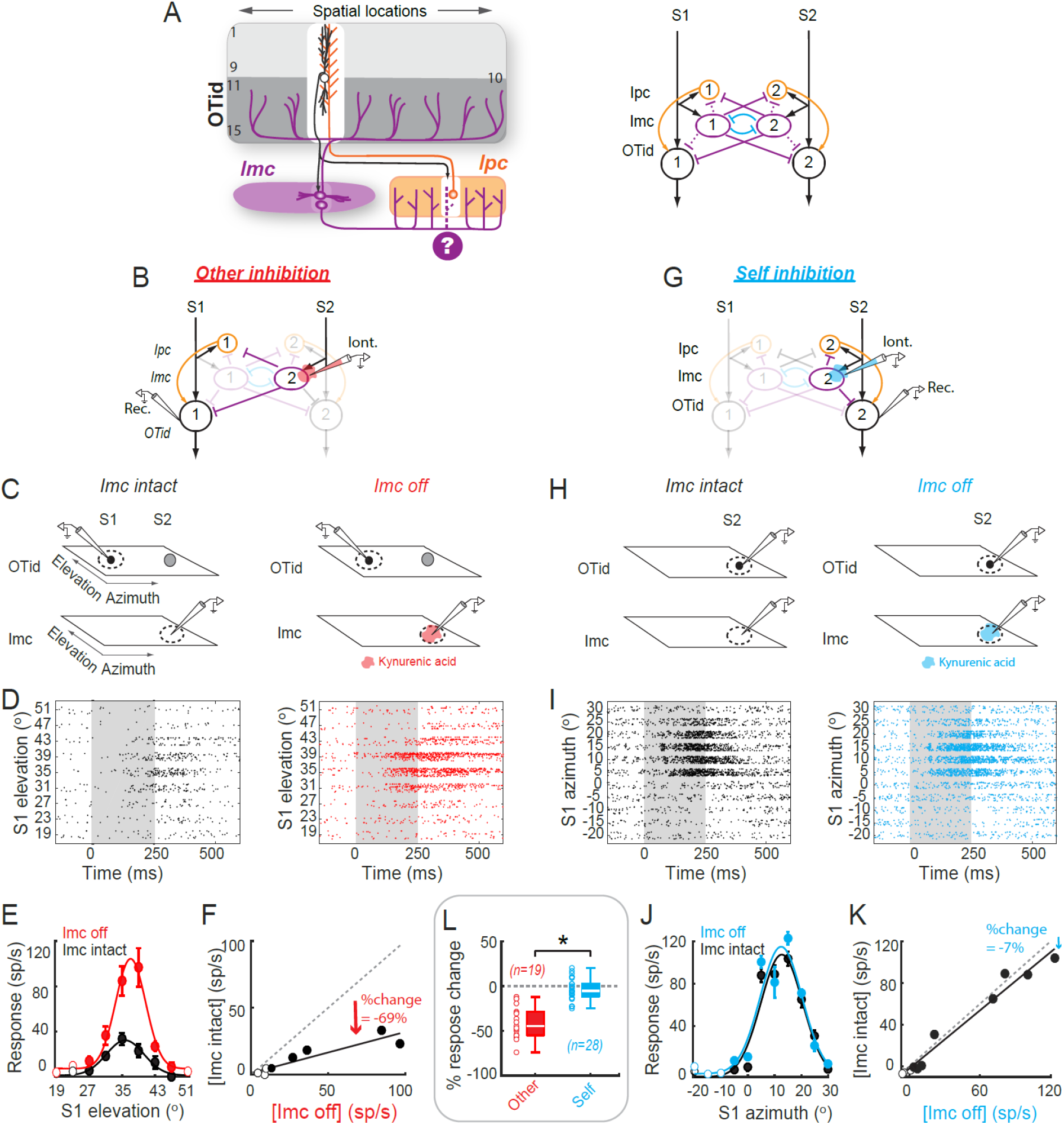
Barn owl midbrain selection network contains functional donut-like inhibitory motif. **(A)** <u>Left:</u>Schematic of avian (barn owl) midbrain selection network; modified from Fig. 1D; same conventions. Columns across layers of OT tissue, from left to right, encode individual locations in space (here, azimuth) topographically^51^. White column in OT space map: Inhibitory projections from Imc neuron that impinge broadly across OTid thought to spare this portion of OT which provides input to that Imc neuron (absence of purple projections here; white column surrounds the black OT10 neuron providing input^31^). Purple ‘?’: unknown if congruent portion of Ipc space map (white column) is also spared of Imc projections (dashed purple projections). <u>Right</u>: Network model showing the OT-Imc-Ipc circuit; conventions as in Fig. 2A. Dashed purple lines: Unknown if these ‘self’-inhibition connections exist functionally; in case of Imc→ Ipc, anatomical evidence is also lacking. **(B-F)** Measurement of the strength of net “other” inhibition from Imc → OTid with paired recordings in barn owl Imc and OTid. “Net” indicates the combined inhibition due to both the direct (Imc→ OTid) and indirect (Imc→ Ipc→ OTid) pathways (text). “Other” indicates that the OTid neuron encodes for (distant) spatial locations outside Imc neuron’s RF. (B) Experimental setup. Iontophoresis and recording electrode (Iont.) in the portion of Imc (encoding for stimulus S2); recording electrode (Rec.) in the portion of OTid encoding for distant location (and stimulus S1). Neurons and connections not immediately relevant to current experiment are shown ghosted-in. (C) Schematic of OTid and Imc space maps (quadrilaterals) showing RFs of neurons being recorded (dotted ovals) and stimulus protocol (black filled dot – S1; gray filled dot – S2). S1 and S2 are looming visual stimuli of fixed contrast but different loom speeds (strength ^49^); S1 = 9.6 °/s, S2 = 19.2°/s (Methods). (D) Raster responses of example OTid neuron to paired stimulus protocol in C, in the Imc-intact condition (left column; black data) and Imc-off condition (right column – red data). Imc inactivation by (reversible) iontophoresis of kynurenic acid (a pan-glutamate receptor blocker^32^; Methods). Gray shading: stimulus duration (E) Response firing rates of this OTid (computed from rasters in D; spike count window = 100-350ms); mean ± s.e.m Lines: best Gaussian fits. Filled dots: responses to S1 at locations inside the OTid RF; open circles, outside RF (Methods). (F) Scatter plot showing OTid responses in Imc-intact vs. Imc-off conditions. Line: Best fit straight line; slope = 0.31, r^2^ =0.76. % change in responses = 100* (responses in Imc intact condition – responses in Imc off condition)/responses in Imc off condition = 100*(slope-1), directly estimates the strength of inhibition at this OTid neuron due to Imc (here, −69%; 100*(0.31-1); Methods). **(G-K)** Measurement of the strength of net “self” inhibition from Imc → OTid with paired recordings in Imc and OTid. (conventions as in B-F). (G, H) OTid neuron is spatially aligned with Imc neuron (both encode overlapping locations); distance between OTid and Imc RF centers = 1.5°. (K) Line: Best fit straight line; slope = 0.93, R^2^ =0.95. % change in responses directly estimates the strength of suppression at this OTid neuron due to Imc (here, −7%; Methods). **(L)** Population summary of strength of net “other” inhibition (red; n=19 Imc-OTid pairs), and strength of net “self” inhibition (blue; n=28 pairs) from Imc→ OTid. Average distance between OTid and Imc RF centers in “other” experiments = 26.8° +/- 2.3°; in “self” experiments = 2.9° +/- 0.7 °. p<1e-5 (red vs. blue), p<1e-5 (red vs 0), p=0.12 (blue vs. 0), paired t-tests with HBMC correction (Methods). **See also Fig. S2.**

The eight circuit models that arise from different combinations of these three proposed circuit motifs are illustrated in Figure 2A. We next examined the ability of each model to produce categorical neural response profiles. We did so by measuring the responses of a designated output neuron in each model (Fig. 2A, ‘OTid’ neuron 1) to a classic two-stimulus strength-morphing protocol^2,6,8^ (Fig. 2B). This protocol was identical to that used in past experimental work^8^, and involved the simultaneous presentation of two stimuli (S1 and S2) at distant spatial locations (or to distinct ‘channels’ in the model). The relative strength of the S1 and S2 (controlled by their loom speed) was systematically varied, resulting in two well-defined stimulus strength-dependent categories^2,6,8^: S1> S2 and S1<S2; Fig. 2B; grey line). Model neurons in the circuit were simulated with noisy, sigmoidal input-output functions (30 repetitions per ‘neuron’, n=50 ‘neurons’; Methods). The values of the parameters of these sigmoids, as well as the value of response fano factor used in the models (= 6; Methods), were obtained from a series of previous experimental (electrophysiological) measurements of input-output functions in the barn owl midbrain network^9,28,30,44,48,49^.

To quantify the strength of categorization of the resulting response profiles, we used a generalized version of the classic categorization index used in the literature ^6–8^. This index, defined as the average difference in responses between categories divided by the average difference in responses within categories, is insensitive to simple multiplicative scaling of the responses (Fig. S1AB), and quantifies how ‘step-like’ the response profiles are. Here, we modified it to define a generalized categorization index, ‘*CatI*’, which, in addition, takes into account neural response variability as well. *CatI* is defined as the average difference in discriminability (d’) between categories divided by the average difference within categories (Methods; Fig. 1A, right panel; Fig. S1AB). We computed *CatI* for the strength-dependent response profiles from each of the eight models (Fig. 2A), and compared them statistically (Fig. 2D; ANOVA with HBMC correction; Methods).

Our simulation results revealed that feedback inhibition between the competing channels, reflecting inhibition between Imc neurons encoding for S1 and S2 in the avian midbrain network, had no significant effect on *CatI* (Fig. 2C-inset and 2D – green vs. grey; p =0.99). This result was largely independent of the strength of feedback inhibition (Fig. S1D). Similarly, recurrent excitation within each channel, reflecting Ipc amplification of OTid activity in the avian midbrain network ^25,27^, had no significant effect on *CatI* (Fig 2C-inset and 2D-orange vs. grey). This result as well was largely independent of the strength of recurrent amplification (Fig. S1E). Furthermore, the combination of feedback inhibition and recurrent excitation also had no significant effect on *CatI* (Fig. 2D-brown). These results held true also when we simulated an alternative implementation of recurrent excitation, one that has been used more generally in modeling work of decision-making^40,42^. In this variant, called just ‘recurrence’ (as opposed to ‘Ipc recurrence’), the strength of amplification (Fig. S1F, top-left, orange arrow), is not regulated by competitive inhibition (from ‘Imc’; compare Fig. S1F, top-left to Fig. 2A, left column – bottom). Introduction of such recurrence, either by itself, or in conjunction with feedback inhibition, did not produce categorical responses. (Fig. S1GH).

By contrast, however, a donut-like pattern of inhibition from Imc to OTid in the model substantially boosted *CatI* of the response profiles (Fig. 2C-inset and 2D–purple vs. grey; p = 5.98e-8). The magnitude of improvement was inversely related to the strength of ‘self’-inhibition, reaching the maximum when self-inhibition was zero, i.e., when the pattern of inhibition was fully donut-like (Fig. S1I; *CatI*: ρ*=-0.81*, *p = 2.6 e-3,* Pearson correlation test). Whereas the presence of either or both recurrent excitation and feedback inhibition enhanced the impact of the donut-like motif (Fig. 2CD: blue vs. purple, red vs. purple; p< 6.023 e-8 in both cases), without the donut-like motif, they were nearly ineffective, either individually or together, at signaling the strongest stimulus categorically (Fig. 2CD: green, orange, brown). The magnitude of response variability (fano factor) in responses did not alter these findings (Fig. S1J).

Finally, we explored whether varying the values of various parameters in models not containing the donut-like motif, but containing feedback inhibition and/or recurrent amplification, might allow them as well to achieve categorical responses. We varied several key parameters (including slopes of the input-output functions), and simulated 324 circuit models representing different combinations of parameter values (Fig. S1K). We found that these models were still markedly less effective (at best, half as effective) at generating categorical responses compared to the model containing just the donut-like motif (Fig. S1L).

Donut-like inhibition, therefore, emerged as the most powerful single circuit motif (among the three proposed in the literature) for producing categorical neural responses (Fig. 2D).

### Functional pattern of competitive inhibition in the owl midbrain selection network is donut-like

To examine the potential role of donut-like inhibition in controlling categorical neural responses in the avian midbrain, we first investigated experimentally whether a donut-like inhibitory motif even operates in the owl midbrain selection network. As we pointed out above, anatomical tracing studies^31^ have indicated that the direct projections from Imc neurons to the OT spare the portion of the OT providing input to Imc, supporting a donut-like motif in the *direct* inhibitory pathway (Fig. 3A; white column – highlighting absence of purple projections in portion of OT containing black neuron), although this not been established functionally. Crucially, however, whether or not the *indirect* inhibitory pathway involving the Ipc also exhibits the donut-like motif is unknown (Fig. 3A, purple ‘?’ under dashed projections and white column in Ipc). This is critical because the indirect pathway is known to be the dominant route of inhibition from Imc to OTid^31^: the majority of Imc projections target the Ipc^31^ (rather than the OTid), and Ipc provides substantial amplification of OTid responses to single stimuli^25^.

To determine whether the Imc-Ipc-OT circuit implements a functional donut-like pattern of competitive inhibition, we measured experimentally the strength of the net inhibition delivered by Imc neurons onto OTid neurons, due to both the direct as well as the indirect pathways. Specifically, we compared the strength of net inhibition from Imc neurons onto ‘misaligned’ OTid neurons encoding for stimuli at distant, non-matched azimuthal locations (‘other’-inhibition; Fig. 3B), and separately, that onto ‘aligned’ OTid neurons encoding for overlapping locations (‘self’-inhibition; Fig. 3G). We did so by making dual extracellular recordings in the barn owl OTid and Imc (Methods; Fig. 3BG).

To measure the strength of net ‘other’-inhibition, we first recorded the responses of OTid neurons to a stimulus (S1) inside the receptive field (RF; Fig. 3B, C-left; Methods) while simultaneously presenting a competing stimulus outside the RF (S2; at a distant azimuthal location from S1; Methods). We then repeated this measurement after focally (and reversibly) inactivating the portion of Imc encoding S2 (spatially mismatched with the OTid recording site; Fig. 3C-right), and compared the responses. Inactivation was achieved by iontophoresing the pan-glutamate receptor blocker, kynurenic acid (Methods).

Responses to S1 in the OTid are known to be divisively suppressed by a distant S2 (by an amount depending on their relative strength)^49,50^, and this suppression is known to be abolished upon focally inactivating the portion of Imc representing S2 ^32,33^. Therefore, any increase in OTid responses to the paired presentation of S1 and S2 following Imc inactivation would represent other-inhibition provided by Imc. We quantified the strength of this net other-inhibition as % change in OTid responses: 100*(responses in Imc intact condition – responses in Imc off condition)/responses in Imc off condition (Methods; no change in OTid responses would indicate zero competitive suppression by Imc onto that OTid neuron.)

We found that Imc neurons exerted strong inhibition onto OTid neurons encoding for distant, non-matched spatial azimuths (Fig. 3F, L-red; mean strength = - 40.47 % +/- 17.70 %, n==19 paired neurons; p=9.43e-9, t-test against 0; mean distance between centers = 26.74 °). We verified that the results were specifically due to Imc inactivation by observing that OTid responses to paired S1 and S2 returned to pre-drug levels after recovery from iontophoresis (measured 15 min after the drug was turned off; Fig. S2A; Methods). Indeed, the suppression provided by Imc accounted for nearly all the suppression exerted by the stimulus S2 (Fig. S2B). In these experiments, Imc was inactivated effectively (median = 95%, 95% CI of median = [87%, 103%]; p = 3.8e-6, sign test, n=19; Fig. S2C).

Next, to measure the strength of net ‘self’-inhibition, we recorded the responses of OTid neurons to a single stimulus (S1) presented inside the RF (Fig. 3G, H-left; Methods). We then repeated this measurement after focally (and reversibly) inactivating the portion of Imc also encoding for S1 (spatially matched with the OTid recording site; Fig. 3H-right; Methods), and compared the responses.

Following a similar argument as above, any increase in OTid responses between the Imc-intact (Fig. 3I-left; 3J-black) and the Imc-inactivated (Fig. 3I-right; 3J-blue) conditions would represent self-inhibition provided by Imc. We quantified the strength of this net self-inhibition also as % change in OTid responses: 100* (responses in Imc intact condition – responses in Imc off condition)/responses in Imc off condition (Methods).

We found that Imc neurons exerted no significant inhibition onto OTid neurons encoding for overlapping spatial locations (Fig. 3K, L-blue; mean strength = −3.7 %, s.d. = 12.2%, n = 28 neuron pairs; p = 0.12, t-test against 0; mean distance between centers = 2.86 °). We verified that these results were not due to ineffectiveness of iontophoresis by observing that the suppression of Imc responses by kynurenic acid was substantial (Fig. S2C), and not distinguishable from that in the ‘other’ case (Fig. S2C; p = 0.68, ranksum test, Imc suppression by drug in ‘self’ vs. ‘other’ cases). Therefore, strength of net self-inhibition in the OTid was substantially weaker than the strength of net other-inhibition (Fig. 3L, red vs. blue; p = 7.8e-11, two sample t-test with HBMC correction; Methods). Additionally, the average strength of net self-inhibition from Imc to OTid was not significantly different from zero (Fig. 3L, blue; p = 0.12, t-test with HBMC correction; Methods).

Together, these findings demonstrated the presence of a net functional donut-like pattern of competitive inhibition implemented by Imc across the OTid (azimuthal) space map; operating necessarily along both the direct and indirect pathways. Notably, they attest to the presence of a functional donut-hole of inhibition in Imc→OTid projections, as well as in Imc→ Ipc projections.

### Donut-like inhibition in the avian midbrain selection network is multi-holed

Imc neurons are strikingly asymmetric in their encoding of elevational versus azimuthal space. The majority (67%) exhibit RFs with multiple discrete firing fields (“lobes”) distributed along the elevation, but not the azimuth (multilobed RFs exhibit up to 3 lobes along the elevation^52^; Fig. 4A,G-left panels). This unusual RF structure has been shown to be essential for Imc to achieve selection at all possible pairs of spatial locations in the face of scarcity of its neurons, and it does so using a combinatorially optimized inhibitory strategy^52^.

**Fig. 4.**
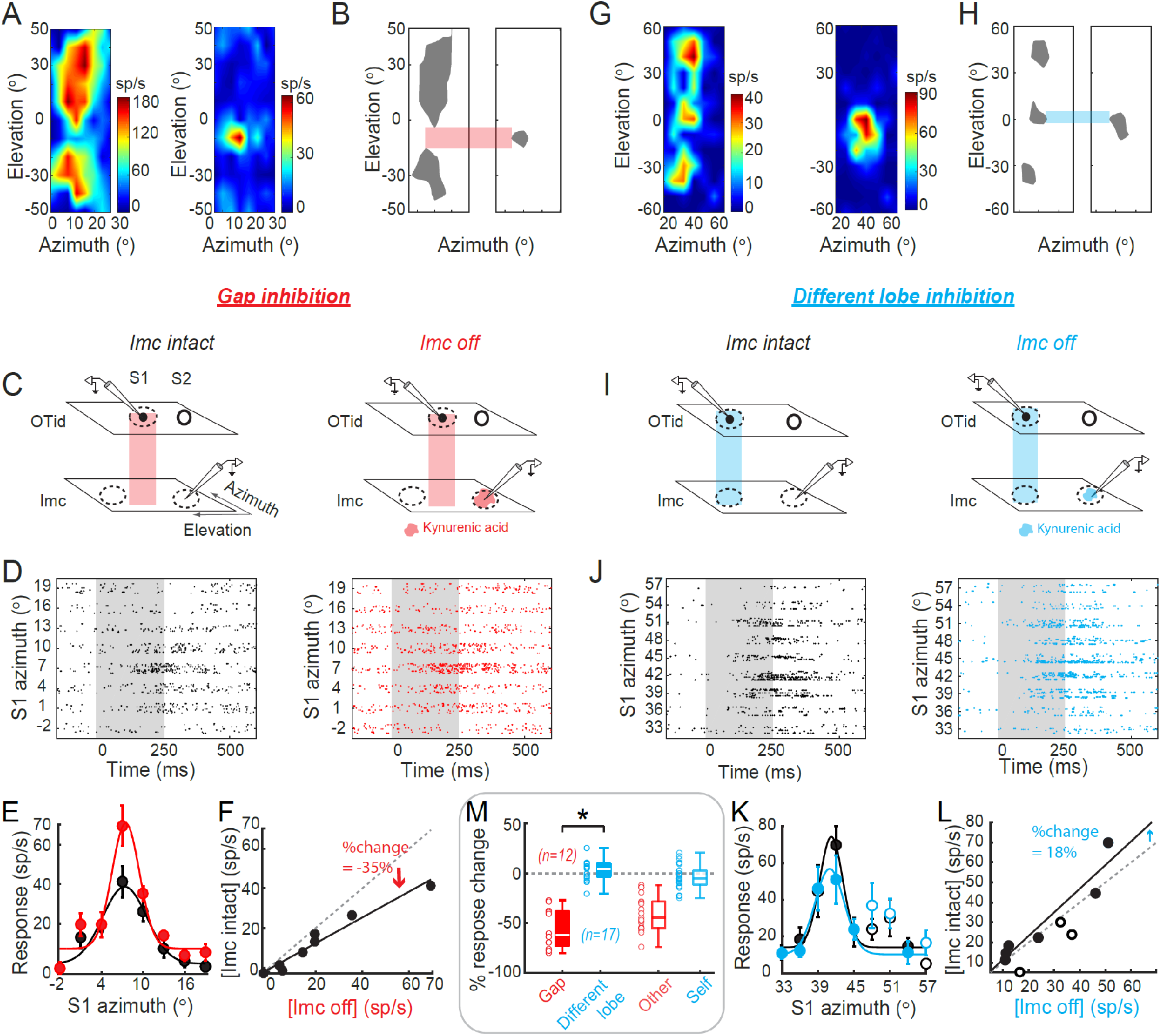
Barn owl midbrain selection network contains multi-holed donut-like inhibitory motif operating across 2-D sensory space (azimuth x elevation). **(A-F)** Measurement of the strength of net “gap” inhibition from Imc → OTid with paired recordings in owl Imc and OTid. (A) Spatial receptive fields (RFs) of an Imc neuron (left) and an OTid neuron (right) from a paired Imc-OTid recording experiment. Imc RF is two-lobed^52^(text; Fig. S3B-D; Methods). OTid RF lies in the gap between the lobes of the Imc RF. (B) Binarized versions of RFs in A, at 60% max. firing rate in each case. Red horizontal bar: highlights the relative position of OTid RF to Imc RF lobes. (C-F) Conventions as in Fig. 3C-F. C, Red vertical bar indicates that OTid RF is in the gap between Imc RF lobes. F, Line: Best fit straight line; slope = 0.65, R^2^ =0.95. % change in responses directly estimates the strength of inhibition at this OTid neuron due to Imc (here, −35%; Methods; conventions as in Fig. 3F). **(G-L)** Measurement of the strength of net “different lobe” inhibition from Imc → OTid with paired recordings in Imc and OTid. Conventions as in A-F. (G, H) Three-lobed spatial RF of an Imc neuron (left; Fig. S3E-G; Methods). RF of OTid neuron overlaps (blue bar) with one of the lobes of Imc neuron’s RF. (L) Line: Best fit straight line; slope = 1.18, R^2^= 0.9. % change in responses directly estimates the strength of inhibition at this OTid neuron due to Imc (here, 18%; Methods; conventions as in Fig. 3F). **(M)** Population summary of strength of net “gap” inhibition (red; n=12), and strength of net “different lobe” inhibition (blue; n=17) from Imc→ OTid. Open red and blue box plots: reproduced from Fig. 3L. p<1e-5 (red vs. 0), p=0.35 (blue vs. 0); p=<1e-5 (red vs blue), paired t-tests with HBMC correction. **See also Fig. S3.**

A direct consequence of multilobed encoding of elevational space is that there are gaps between the lobes of an Imc RF, constituting locations that are outside that neuron’s RF (Fig. 4AB-left panels; light red bar). If the donut-like inhibitory motif is to operate generally in the avian midbrain network to support selection along not only azimuthal locations, but also along elevational locations, then the spatial pattern of inhibition must respect the following strict conditions. A multilobe Imc neuron activated by a stimulus within one of its RF lobes must send strong competitive inhibition to OTid neurons encoding locations outside all of its RF lobes, and specifically, to neurons encoding locations in the gaps between RF lobes (strong ‘gap’-inhibition). By contrast, it should send weak or no inhibition to OTid neurons encoding locations within any of its RF lobes, and specifically, within its other RF lobes (weak ‘different-lobe’ inhibition). Together, these predict a multi-holed donut-like pattern of net inhibition from Imc to OTid (Fig. S3A).

To test experimentally if this strict requirement holds true in the owl midbrain selection network, we again made dual extracellular recordings in the OTid and Imc, and measured directly the strength of gap-inhibition, and separately, the strength of different-lobe inhibition from Imc neurons onto OTid. We first recorded the responses of an Imc neuron, mapped out its spatial RF, and applied previously published analytical approaches to determine if it was a multilobed RF (two-lobed RF in Fig. 4AB – left panels and Fig. S3B-D; three-lobed RF in Fig. 4GH – left panels and Fig. S3E-G^52^; Methods). If so, we next measured the strength of gap inhibition by positioning a second electrode in the OTid such that the spatial RF of the OTid neuron was centered within the gap between Imc RF lobes (Fig. 4AB – right panels and Fig. 4C-left, light red bar). We recorded the responses of the OTid neuron to a stimulus inside its RF (S1; Fig. 4C-left; Methods) while simultaneously presenting a competing stimulus (S2) at a distant location along the elevational axis such that S2 was within a lobe of the RF of the Imc neuron (Fig. 4C-left – S1 in the Imc RF gap denoted by light red bar, and S2 within an Imc RF lobe).

Alternatively, to measure the strength of different-lobe inhibition, we positioned the OTid electrode such that the spatial RF of the recorded OTid neuron overlapped one of the lobes of the Imc neuron’s RF (Fig. 4GH – right panels and Fig. 4I – left, light blue bar). We recorded the responses of the OTid neuron to a stimulus inside its RF (S1; Fig. 4I-left; Methods) while simultaneously presenting a competing stimulus (S2) at a distant location along the elevational axis such that S2 was within a different lobe of the Imc neuron’s RF than S1 (Fig. 4I-left – S1 within one Imc RF lobe and S2 within different one – light blue bar).

In both cases, we compared OTid responses when Imc was intact (Fig. 4CI-left panels) versus when the portion of Imc encoding S2 was focally and reversibly inactivated (Fig. 4CI-right panels). As before, any observed response increases directly estimated, respectively, the (net) strengths of gap-inhibition or different-lobe inhibition exerted by an Imc neuron onto the OTid space map (Methods).

We found that Imc neurons exerted strong gap-inhibition (Fig. 4D-F, M-red), but weak different-lobe inhibition onto OTid (Fig. 4J-L, M-blue). The mean strength of gap-inhibition (along elevation) was strong at −55.88% (Fig. 4M-red, s.d. = 20%, n=12 neuron pairs; p = 1.02 e-6, t-test against 0 with HBMC correction; Fig. S3IJ-red; Methods). The mean strength of different-lobe inhibition was very weak at 2.43 % (Fig. 4M-blue, s.d. = 10.3 %, n=17 neuron pairs; p = 0.35, t-test against 0 with HBMC correction; Fig. S3IJ-blue; Methods), and was not distinguishable from the average self-inhibition (Fig. 4M-blue vs. open blue; p-value = 0.09; two sample test with HMBC correction). Imc was inactivated effectively in both sets of experiments (median = 98.2%, 95% CI of median = [96.46% 99.99%]; Fig. S3K).

Taken together, these results demonstrated that the Imc implements precisely organized, multi-holed donut-like patterns of net inhibition onto the OTid space map (azimuth and elevation), operating along both the direct and indirect pathways. Each Imc neuron’s net inhibitory action in the OTid creates a spatial pattern complementary to its (multilobed) RF structure (Fig. S3A).

### Donut-like inhibitory motif is necessary for categorization by avian midbrain selection network

Considering the complexity of the multi-holed donut-like connectivity between Imc (and Ipc) and OT (Fig. 5A), we next asked if this served a functional purpose. Specifically, we investigated whether this motif was necessary for the categorical signaling of the strongest stimulus by the OTid. To test this, we needed to test the impact of causally disrupting the donut-like pattern of inhibition from an Imc neuron to OTid on categorical signaling in the OTid. In other words, we needed to selectively introduce self-inhibition onto the spatially aligned OTid neuron, thereby ‘filling-in’ the donut hole (Fig. 5B, gold projections). However, introducing self-inhibition experimentally by activating Imc → Ipc projections (indirect pathway) or Imc → OTid projections (direct pathway) between spatially aligned neuron-pairs (Fig. 5B, gold projections) is not feasible because such projections either do not exist or are functionally inactive (combined self-inhibition at most OTid neurons is weak or zero; Figs. 3L and 4M).

**Fig. 5.**
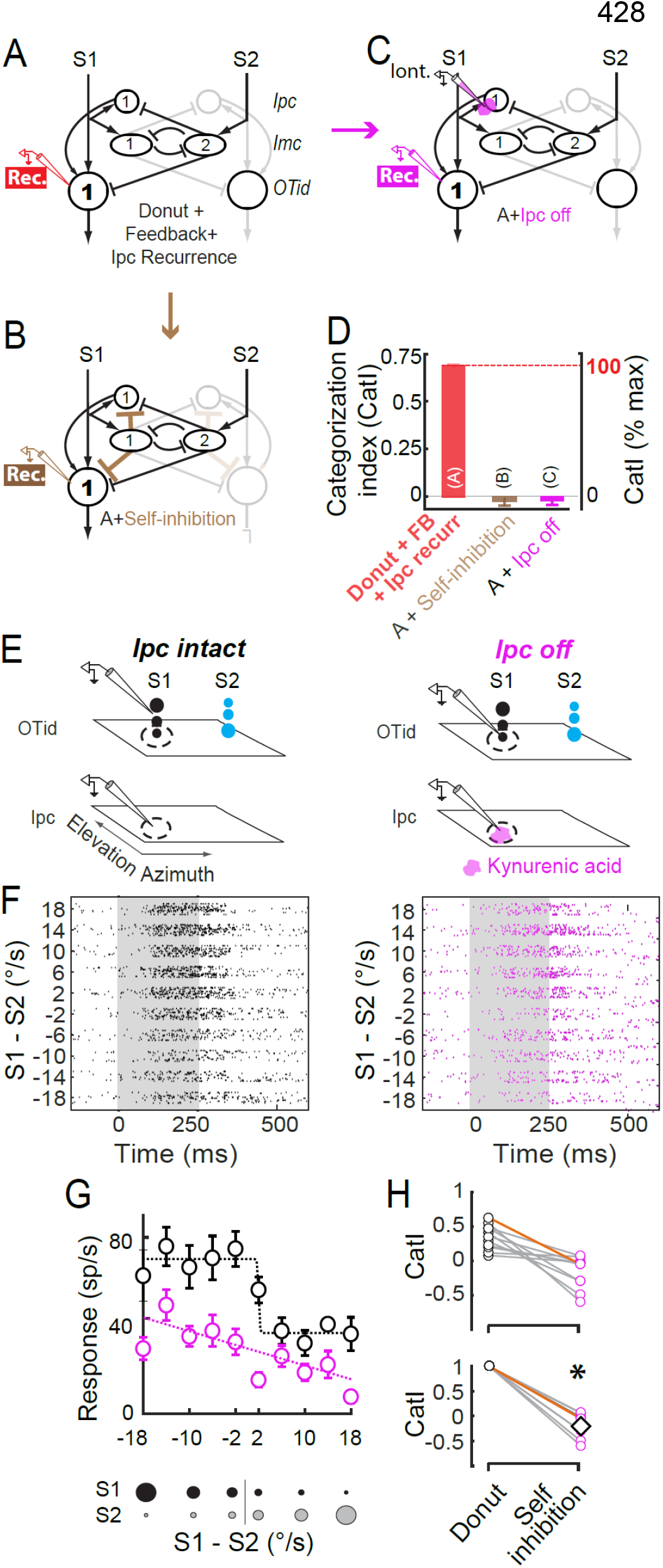
Donut-like inhibitory motif is required for categorical signaling of the strongest stimulus by the barn owl midbrain selection network. **(A)** Network model schematizing the OT-Imc-Ipc circuit with donut-like inhibition in the barn owl midbrain. ‘Rec’: Recording at OTid neuron 1. **(B)** Network model schematizing the desired experimental manipulation of disrupting donut-like inhibition (i.e., filling-in the donut-hole): introduction of self-inhibition (brown connections) in the direct pathway between Imc and OTid, a s well as in the powerful indirect pathway through Ipc. However, this is infeasible experimentally (Figs. 3 and 4; text). **(C)** Network model schematizing the proposed equivalent experimental manipulation: focal inactivation of Ipc neuron in channel 1, mimicking the effect of introducing self-inhibition onto Ipc neuron 1, while recording from aligned OTid neuron 1. ‘Iont’: Iontophoresis of kynurenic acid (magenta blob) onto Ipc neuron; ‘Rec’: Recording OTid neuron. **(D)** Modeling results showing equivalence between desired (B) and proposed (C) manipulations. Plots of CatI computed from response profiles simulated from models in A-C; conventions as in Fig. 2AB. Manipulations in both B and C cause abolishment of categorization. See also Fig. S4A-C showing, additionally, that C is not equivalent to removal of recurrent amplification. (Note: model in A is identical to model in Fig. 2A, right column-bottom; model in B is identical to model in 2B, middle column-third from top). **(E)** Paired recordings in OTid and Ipc such that RF of OTid neuron overlaps with that of Ipc neuron. Conventions as in Fig. 3C. Stimulus protocol used is the same two-stimulus morphing protocol used in model simulations in Fig. 2B-D; the relative strength between S1 and S2 was systematically varied. **(F)** <u>Left</u>: OTid response rasters in the Ipc-intact condition. <u>Right</u>: OTid response rasters in the Ipc-off condition. Distance between OTid and Ipc RF centers = 5°. **(G)** OTid response firing rates, computed from D over the 100-400ms time window. Black: Ipc-intact condition, magenta: Ipc-off condition. Dashed lines: best fitting sigmoid or straight line to data, chosen based on AIC criterion. Black: AIC (sigmoid) = 29.07, AIC (line) = 40.92; magenta: AIC (sigmoid) = 37.75, AIC (line) = 34.67. **(H)** Population summary of effect of Ipc inactivation on CatI; n=11 neuron pairs; data in orange are from example neuron pair in F-G. Top panel: measured values. Bottom panel: Data in top panel replotted after normalizing to Ipc-intact values. Diamond: average of magenta data;*: p <1e-4, ranksum tests against 1 with HBMC correction. **See also Fig. S4.**

To resolve this apparent conundrum, we stepped back and considered the specialized anatomical connectivity of the avian midbrain selection network (Fig. 5A). We observed that in a competitive setting (i.e., when S1 and S2 are both present), the ‘amplifier’ Ipc neurons corresponding to each stimulus are not free to amplify OTid activity solely based on that stimulus’s strength. For instance, whether Ipc neurons encoding for S1 amplify OTid responses to S1 at all, and if so, to what extent, are controlled by the strength of competitive inhibition due to S2 delivered by Imc neurons in S2’s channel (Fig. 5A, oval ‘Imc’ neuron #2). In other words, in a competitive setting, the ‘amplifier’ Ipc neurons operate wholly under the powerful control of the inhibitory Imc neurons.

This observation led us to reason that we could achieve the goal of experimentally introducing self-inhibition onto OTid neurons not by activating the (non-existent) self-projections from Imc to the aligned Ipc neuron in the dominant indirect pathway (Fig. 5B), but rather by mimicking the consequence of doing so, namely, by focally suppressing the evoked output of the aligned Ipc neuron (Fig. 5C, pink, drug iontophoresis onto aligned Ipc).

To test if this logic was sound, we first simulated this latter model (Fig. 5C), and found that focal Ipc-inactivation in the model abolished categorical signaling by OTid (Fig. 5D, pink vs. red), and this was indistinguishable from the effect of introducing self-inhibition (Fig. 5D, gold versus red; simulation of 5B). This established the effectiveness of the proposed manipulation. Notably, this manipulation in the avian midbrain circuit during stimulus competition was akin to filling-in the donut hole, rather than to simply silencing recurrent amplification: simulating the silencing of just the recurrent amplification in a circuit model in which recurrence was not under the control of powerful competitive inhibition did not produce a significant drop in categorical signaling (Fig. S4A-C, blue vs. red-gray). In other words, the result of Ipc inactivation (5C, D) was nearly identical to that of introducing self-inhibition (5B, D), but very different from that of silencing recurrent amplification (Fig. S4B, C). These model simulations established that in the avian midbrain selection network, focal Ipc inactivation in a competitive setting (Fig. 5C) was functionally equivalent to the desired causal manipulation of filling-in the ‘hole’ in the donut-like pattern of inhibition.

Using this insight, we proceeded to test experimentally, the functional consequence of filling-in the donut hole on categorization by barn owl OTid neurons. We focally (and reversibly) inactivated Ipc neurons by the iontophoresis of the pan-glutamate receptor blocker, kynurenic acid (Fig. 5E-right, pink blob, ‘Iont.’; Methods), while simultaneously recording the responses of spatially matched OTid neurons (‘Rec.’). OTid responses were measured to the same two-stimulus morphing protocol used in our modeling (Fig. 5E left vs. right; protocol shown in Fig. 5G–bottom, same as Fig. 2C; Methods), as well as in previous experimental studies of categorical signaling in the OTid^8,22^.

We found that disrupting the donut-like inhibitory motif caused a substantial reduction in categorical signaling in the OTid (Fig. 5FG: example neuron pair). Across the population of tested neuron pairs, categorization was nearly abolished by this experimental perturbation, with the median reduction in *CatI* of 104.7% (Fig. 5H; n=11 neuron pairs; *CatI*: 95% CI of median = [91.52%, 117.92%]; p = 2.55 e-5, ranksum test against 1 with HBMC correction). These results reinforce the dominance of the indirect Imc-Ipc-OT pathway over the direct Imc-OT pathway (the donut-like inhibition there remained intact in these experiments). Thus, the midbrain spatial selection network not only contains a specialized donut-like inhibitory circuit motif, but also critically depends on it for categorical signaling by OTid.

## DISCUSSION

Our results discover the donut-like inhibitory motif as a powerful, identifiable circuit mechanism for generating categorical neural selection boundaries, able to convert even linear response profiles to categorical ones (Fig. 2D-inset, 2D – orange/brown to purple; Fig. 5GH – pink vs. black data).

### Superiority over feedback inhibition and recurrent excitation motifs for generating categorical neural responses

Contrary to prior proposals^42,44^, our results showed that feedback inhibition and recurrent amplification, by themselves, are not effective at producing categorical response profiles, nor is their combination (Fig. 2CD). This was true independently of the strength of feedback inhibition (Fig. S1D), the strength of recurrent amplification (Fig. S1E), the specific implementation of recurrent amplification (Fig. S1F-H), the magnitude of response noise (Fig. S1J), and over a range of values of key parameters of the models including the slopes of the single-stimulus input-output functions (or neuronal ‘activation’ functions; Fig. S1KL). (The latter point is consistent with previous work showing that the steepness of input-output functions of neurons can be uncorrelated with whether the selection signal is categorical^9^.) Although not effective for producing categorical responses, feedback inhibition and recurrent amplification have been linked to other important functions related to selection, namely, the implementation of a flexible selection boundary (feedback inhibition^13,44,53^), and evidence accumulation (recurrent amplification^39,42^). Selection circuits may, therefore, need to include these motifs for other reasons than categorization. If present as well, they heighten the efficacy of the donut-like inhibitory motif in producing categorical response profiles (Fig. 2D and S1H), effectively helping implement attractor dynamics for flexible and categorical decision-making^2,40,44^.

A viable alternative to donut-like inhibition for categorization are highly recurrent, non-structured networks, which have been shown in modeling to be capable of generating categorical outputs from a multiplexed representation of inputs ^54,55^. However, because it is difficult to extract specific, experimentally testable neural circuit mechanisms from the opaque connectivity diagrams of recurrent networks, and in light of the recently reported counterpoint to mixed-selectivity descriptions^10^, we focused, here, on structured circuit mechanisms, and identified donut-like inhibition as being highly effective for categorizing inputs.

### Superiority over the normalization model for generating categorical neural responses

The structured donut-like organization of inhibition stands in contrast to another computational mechanism that has been invoked in the decision-making literature, namely divisive normalization^46,56–58^. This involves inhibitory elements that pool the drive from all the active channels (as opposed to receiving selective drive), that inhibit one other, and that deliver pooled inhibition uniformly (rather than in a donut-like manner) to the output elements (Fig. 6A^56,57^). The midbrain spatial selection network (OT-Imc-Ipc), in which categorical neural responses have been reported, does not implement pooled divisive normalization for selection across space: past work has shown that the inhibitory Imc neurons receive selective input^31–33,52^, and our results (Figs. 3-4) show that pattern of inhibitory output across the OT is not uniform (but rather donut-like).

**Fig. 6.**
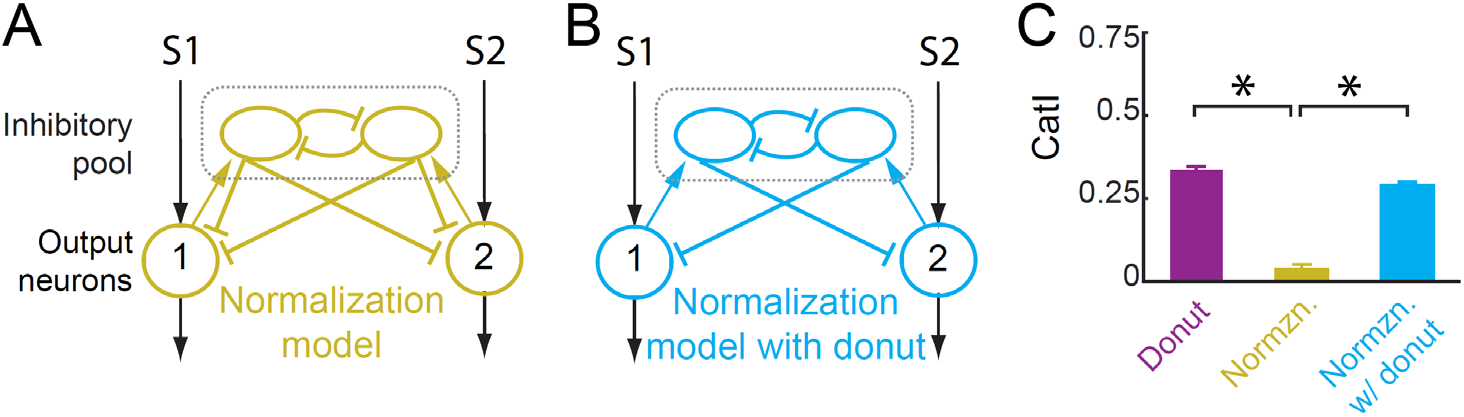
Divisive normalization model is not effective at producing categorical response profiles. **(A)** Schematic of normalization circuit model with pooled inhibition and inhibitory feedback^56,57^. **(B)** Schematic of normalization circuit with self-inhibition removed (i.e., with donut-like inhibition introduced). **(C)** Plot of CatI computed from the responses of neuron 1 in the normalization model (A; gold data) and in the model lacking self-inhibition (B; blue data), to the standard two-stimulus morphing protocol (as in Fig. 2C). For comparison, the CatI values for the model from Figure. 2D that contains the donut-like motif (purple), is reproduced here; p = 5.2e-23 (gold vs. purple), p=8.8e-19 (gold vs. blue), paired t-tests with HBMC correction.

To test more generally whether the divisive normalization model is capable of generating categorical response profiles, we simulated a circuit model of normalization and obtained model neuron responses to the two-stimulus morphing protocol (Fig. 6A). This model yielded a substantially lower *CatI* than the donut-like inhibitory motif (Fig. 6C right panel: mean *CatI* = −0.05 for normalization model compared to 0.331 for donut model in Figure 2A, p = 1.5 e-29, t-test with HMBC correction, gold vs. purple; left panel: p = 1e-12, t-test of gold vs. purple). Thus, the normalization mechanism is not effective for generating categorical responses (consistent with findings from modeling in visual cortex^59^). Indeed, it was the presence of self-inhibition, specifically, that caused this circuit to be ineffective: a modified version of the circuit which did not include self-inhibition (Fig. 6B), did produce categorical responses (Fig. 6C, blue vs. gold data), further attesting to the primacy of the donut-like motif for categorization.

### Alternate implementation of the donut-like inhibitory motif

Both in the barn owl brain and in the various models considered in Figures 2-5, the circuit architectures included feedforward inhibition from inhibitory (‘Imc’) neurons to the output (‘OTid’) neurons, with the donut-like motif instantiated as the absence of feedforward self-inhibition. However, other established models of decision-making do not include feedforward inhibition, but rather only involve inhibition in a ‘reverberant’ path between the competing options ^40^ (Fig. 7A). To test if the implementation method impacted our conclusions, we simulated a version of our circuit model that implemented the donut-like motif only via a reverberant route (Fig. 7A); we note that this implementation, by definition, also includes feedback inhibition between the two channels (Fig. 7A: black neuron 1→ blue neuron 1→ black neuron 2 → blue neuron 2 → black neuron1). For completeness, we simulated this model both without (Fig. 7A) and with recurrent amplification within each channel (Fig. 7B; curved orange arrow). We found that this alternate implementation of the donut-like motif also successfully produced categorical response profiles (Fig. 7CD; *CatI*=0.29 (filled light blue), 0.34 (filled orange)). Nonetheless, the feedforward implementation of the donut-like motif (Fig. 2AB, right panels) offered additional benefits to categorization beyond the purely reverberant implementation in each case (Fig. 7C: filled light blue vs. dark blue; 7D: filled orange vs. red).

**Fig. 7.**
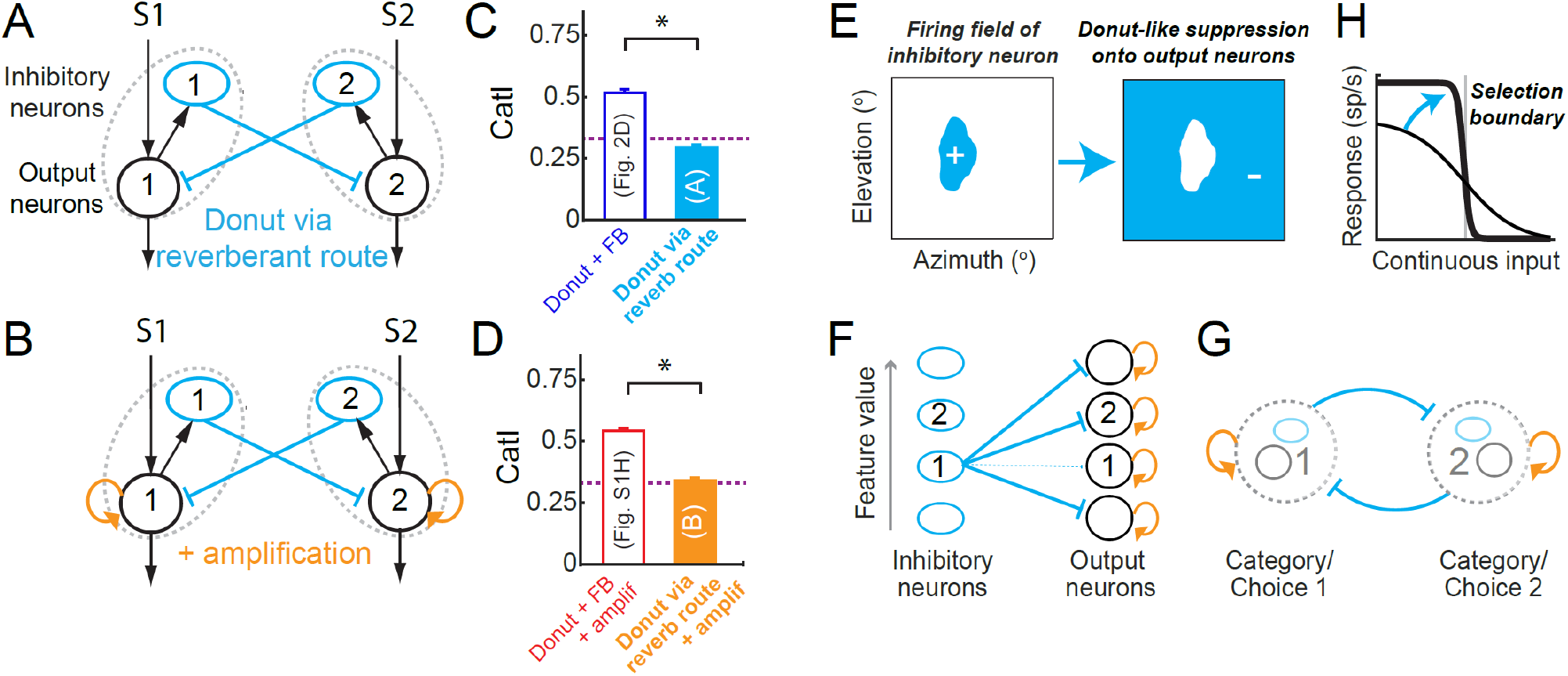
Alternate implementation (and generality) of the donut-like inhibitory motif. **(A,B)** Circuit with implementation of donut-like motif purely via a reverberant route, i.e., in the absence of any feedforward inhibition. (A) Without recurrent amplification. (B) With recurrent amplification; curved orange arrow. Blue ovals: inhibitory neurons, black circles: excitatory/output neurons, dashed gray ovals: populations of neurons representing each stimulus or category. **(C,D)** Plots of CatI computed from the responses of output neuron 1 in circuits in A (C, filled light blue data) and in B (D, filled orange data), to the standard two-stimulus morphing protocol (as in Fig. 2B). For comparison, the CatI values for the corresponding models with feedforward implementation of the donut-like motif are reproduced here (in C: dark blue data from Fig. 2D; in D: red data from Fig. SH). ‘*’: p<0.05. Filled light blue vs. dark blue, p = 1.7e-25; orange vs. red, p = 2.5e-31; paired t-tests with HBMC correction. Purple dashed line: CatI of purple model from Fig. 2A, which has just the feedforward donut-like motif sans feedback; reproduced here from Fig. 2D. **(E-H)** Graphical summary. (E) Donut-like inhibition, i.e., inhibition (-) driven by preferred inputs (+) and delivered to all non-preferred inputs, can be implemented in neural circuits either in a feedforward manner (F) or in a reverberant manner (G), to generate categorical selection boundaries (H). (F,G) Blue ovals: inhibitory neurons, black circles: excitatory/output neurons. F, dashed thin blue arrow indicates absence of inhibitory projection. G, based on Figures 1-4; G is a simplified representation of model in B, in which populations of neurons representing each category or choice (dashed grey circle) mutually inhibit one another. The similarity of this reduced model structure to several previous models of selection highlights the generality of our findings and provides a mechanistic explanation for the ability of those models to produce categorical selection boundaries^40,60–62^; see text. (H) Ability of donut-like motif to convert linear response profiles (thin black) to categorical (thick black) ones

### Generality of donut-like inhibition as mechanism for categorical neural responsess

This study revealed the donut-like inhibitory motif as the engine of categorization in the avian midbrain network for spatial selection, a network conserved across vertebrates^21,31,63,64^. Although the response curves discussed here were obtained using competing looming stimuli, extensive work in the barn owl has shown that categorical response profiles in the owl midbrain are not specific to looming visual stimuli. Rather, they occur no matter what the varying stimulus feature is (stimulus contrast, for instance), and no matter what the sensory modality is (for instance, binaural level of an auditory stimulus)^9,22,28,65^. In other words, categorical neural responses are not idiosyncratic to a specific stimulus feature, but are in fact, general, representing the neural basis of stimulus competition and selection across space in the barn owl^14,66^. Notably, the categorical responses measured in the owl OTid have been shown to be capable of accounting for behavioral deficits in target selection for spatial attention in monkeys following focal perturbation of the homologous SCid ^14^. These observations, together with the fact that the midbrain tecto-isthmic selection network is conserved across all vertebrates^26,31,63,64,67,68^, point to the potential generality of the donut-like motif across the vertebrate midbrain for spatial selection.

Might this motif also generalize to other forms of categorical selection (beyond selection across space), mediated by other cortical and subcortical areas (beyond the midbrain)? If so, how might it generalize, implementation-wise? Below, we examine these two questions in order.

To address the first question of whether or not this motif might generalize (issue of feasibility), we draw upon the extensive literature of computational modeling of categorical decision-making by various cortical (and subcortical) areas ^2, 5–7,11,40,60,61,69,70^. These models successfully account for neural as well as behavioral responses in a wide array of decision-making tasks - perceptual decision-making, delayed-match-to-sample decision-making, sensory categorization, etc. Examination of the circuit architectures in these models reveals that nearly all of them contain a (seemingly hidden) donut-like inhibitory motif^34^. Some include a feedforward implementation ^60,61,69^ (as in Fig. 2 and S1), while others include a reverberant implementation (as in Fig 7) ^40,44, 60–62^. The existence of this motif in these models, however, is not discussed explicitly, and the need for it is unclear. We posit, based on our experimental as well as modeling findings here, that it is this motif that imbues those models with the ability to categorize. Indeed, the one class of models that does not contain the donut-like motif is a normalization-based modeling of decision-making ^71–75^, which we show is ineffective for generating categorical neural response (Fig. 6). Together, these observations support the feasibility of the donut-like motif for producing categorical neural representations across brain areas, animal species and task contexts.

To address the second question of how it might generalize (issues of implementation), we highlight four key characteristics of the donut-like motif in the avian midbrain selection network and discuss plausibly generalizable implementations of each. First, in the midbrain selection network in which the donut-like motif operates, individual stimuli are encoded with neural activity that is proportional to their net priority^21–23^. This readily generalizes to other instances of selection in which stimulus options are encoded with neural activity that is proportional to their net attractiveness or importance: for instance, subjective-value of an option in the case of value-based decision-making^76–78^, degree of membership of a stimulus in the case of perceptual categorization^2,79^, etc. Second, in this network, long-range suppression across spatial locations is the substrate upon which the donut-like inhibitory motif is sculpted, and is implemented by inhibitory neurons with far-reaching projections across the space of stimuli. Alternatively, such inhibition could also be implemented, for instance, in cortical circuits, through long-range excitation contacting local inhibitory neurons. Third, in the midbrain network, the donut-like motif is instantiated via a feedforward implementation. As we saw in Figure 7, an equivalent reverberant implementation, found in many models of selection ^40,80^ is also effective. Fourth, in the midbrain network, the donut-like motif aids spatial selection by operating across a well-organized topographic map of space. However, for many forms of selection, such organized functional maps do not exist, with olfactory categorization being an extreme example^2 81^. In the case of olfactory decision-making, there is some evidence suggesting self-sparing, combinatorial lateral inhibition across mouse glomeruli^81,82^, of the kind found in the owl Imc. In most of these cases, however, a detailed description of the large-scale connectivity diagrams of inhibitory neurons in the corresponding brain areas is yet to be worked out; this study suggests that a search for donut-like inhibitory connectivity might be a fruitful endeavor.

More generally, the donut-like motif described here does not rely on mechanisms of plasticity for its operation. It is able to dynamically generate categorical neural responses from continuously varying inputs, on the fly, and dynamic categorization is critical to various behaviors (value-based decision-making, action selection, spatial attention, etc.). However, conceptually, this motif could also be introduced into a circuit through plasticity mechanisms, thereby aiding in the generation of categorical response profiles that are learned with experience^6,12,83,84^.

### Temporal aspects of selection, and links to behavior

In this study, we were interested, particularly, in mechanisms that could generate categorical profiles of average firing rates (steady state neural activity) as a function of relative stimulus strength, as reported experimentally in the barn owl midbrain^9,22,28,65^. (Similar categorical firing rate profiles have also been reported in multiple other brain areas across species and task configurations^2,5–13^.) Our past modeling work (which forms the basis of this work) has shown that not including precise timing descriptions - for instance, timing of the arrival of excitation versus inhibition, does not impact the ability of these models to capture phenomena at the level of average firing rates^22,44^. Separately, several experimental studies have also reported temporal aspects of decision-making – time courses of neural activity as well as behavioral reaction times. We recall that leading classes of models of selection/decision-making^2,5–7,11,40,60,61,69,70^ contain the donut-like inhibitory motif^34^. Consequently, because those models account for experimentally measured reaction times of animals during the selection tasks (as well as in some cases, response time courses), the donut-like motif is consistent with such temporal aspects and dynamical features of categorical decision-making as well. It will be valuable for future studies (involving behavior) to explore these links explicitly.

The central proposal of this study is that donut-like inhibition underlies categorical neural responses. Whereas neural response profiles underlying decision-making and selection tasks are frequently categorical^2,5–13^, psychometric profiles (of accuracy) are often not. To clarify, whereas the animal or subject must (and does) make a discrete choice on each trial, which results in the task being referred to commonly as a ‘categorical’ decision-making task, behavioral performance, when assessed *as a function of a continuous task parameter*, is often not actually step-like ^17,78,85,86^. It does not exhibit a large, abrupt change across the category boundary, but instead, varies more gradually across it. These two facts, however, are not in conflict: in line with previous findings, neurons can encode information more effectively than the animal as a whole, with behavior being a result of (noisy) aggregation of activity across neurons^87^. Consequently, the central prediction of this study is that causally perturbing donut-like neural inhibition in a relevant brain area during a decision-making task, will cause loss of categorical neural responses, and in turn cause psychometric response curves to become shallower (more gradual) than in the intact condition, independently of their original shape, with selection performance worsening particularly around the selection boundary.

In closing, we propose that the donut-like inhibitory motif may be a critical neural circuit module common to various forms of categorical selection and decision-making. An intriguing open question in this context, even in the avian midbrain selection network, is how the wiring of the exquisitely organized donut-like connectivity is achieved in the brain.

## ACKNOWLEDGEMENTS

This work was supported in part by funding from NIH R01 EY027718. SPM and NRM designed the research and wrote the paper. NRM performed experiments, data analysis and modeling. The authors declare no competing interests.

## METHODS

### Animals

We performed experimental recordings in 7 head-fixed awake adult barn owls viewing a visual screen passively (Tyto alba). Both male and female birds were used; the birds were shared across studies. All procedures for animal care and use were carried out following approval by the Johns Hopkins University Institutional Animal Care and Use Committee, and in accordance with NIH guidelines for the care and use of laboratory animals. Owls were group housed in flight runs within the aviary, each containing up to 6 birds. The light/dark cycle was 12hr/12hr.

### Neurophysiology

Experiments were performed following protocols that have been described previously^32,49^. Briefly, epoxy-coated, high impedance, tungsten microelectrodes (A-M Systems, 250μm, 5-10MΩ at 1 kHz) were used to record single and multi-units extracellularly in the OTid. Multi-barrel glass electrodes (Kation Scientific, Carbostar– 3LT, 0.4-1.2MΩ at 1kHz) filled with kynurenic acid (a competitive inhibitor of ionotropic glutamate receptors; pH 8.5-9 at a concentration of 40mM) were used to record from and inactivate neurons in the Imc and Ipc. Inactivation was performed using micro iontophoresis by ejecting kynurenic acid with an eject current of - 450nA to -500nA; data were collected starting 15 min after drug ejection commenced. A retain current of +15nA was used to prevent leakage of the drug from the tip of the electrodes when drug was not being iontophoresed. Recovery data were measured 15 min after drug ejection was ceased.

### OT, Imc and Ipc targeting

We navigated to the OT (based on well-established methods^51^), and then navigated to the Imc using the OT’s topographic space map as reference. The Imc is an oblong structure that is 2.8mm rostrocaudally and 0.35mm dorsoventrally, appearing as a 700-μm x 350-μm elliptical disk in coronal sections. It lies parallel to the rostrocaudal axis of the OT, located approximately 16 mm ventral to the surface of the brain, and approximately 500 μm medial to the medial-most part of the OT. We targeted the Imc following published methods^32^. Imc targeting has been validated previously using dye injections and lesions^52^. Dorsoventral penetrations through the Imc were made at a medial-leading angle of 5° from the vertical to avoid a major blood vessel in the path to the Imc. The Ipc lies roughly 500-700 um medial to the Imc and its targeting was confirmed based on the neural response characteristics of the neurons (characteristic bursty responses^65^).

### Data collection and spike sorting

Multi-unit spike waveforms were recorded using Tucker Davis Technologies hardware interfaced with MATLAB. The responses of neurons were measured by counting spikes during a time window (typically 100-350 ms) following stimulus onset.

The automated ‘wave-clus’ spike-sorting toolbox was used for spike sorting^88^. We included only those units for analysis for which fewer than 5% of the recorded spikes were within 1.5ms (inter-spike interval; ISI) of each other.

### Model details (Related to Figs. 2, S1, S4, 6 and 7)

#### Input output functions

The input output functions (firing rate *f*, as a function of the saliency, *l*) of the neurons in the model were simulated using sigmoid functions using previously published methods ^44^.

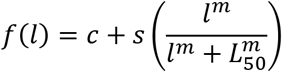

where *c*, is the baseline firing rate of the neuron; *l*, is the saliency parameter of the stimulus (e.g., loom speed of the stimulus, loudness of an auditory stimulus, contrast of the stimulus, speed of a moving stimulus) that can vary continuously over a range; *s*, is the maximum change in the firing rate of the neuron; *L*_50_ is the saliency value at which the neuron’s firing rate changes by 50% of the maximum change; and m, is a parameter that controls the slope of the sigmoid.

#### Excitatory neurons

The excitatory neurons were simulated using the following parameters (which are consistent with the parameters obtained by fitting a sigmoid to response functions of OTid neurons^44^):

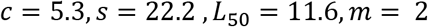

#### Inhibitory neurons

The inhibitory neurons were simulated using the following parameters (which are consistent with the parameters obtained by fitting a sigmoid to response functions of Imc neurons^44^):

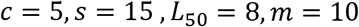

#### Recurrent excitation neurons

The excitatory neurons that provide recurrent amplification in Fig. 2 were simulated using the following parameters (which are consistent with the parameters obtained by fitting a sigmoid to response functions of Ipc neurons^48^):

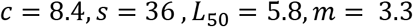

The inhibition sent from the inhibitory neurons onto excitatory neurons is modeled using input and output divisive factors as below using previously published methods^44^.

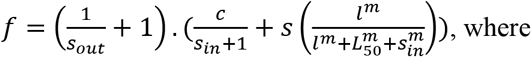

- *s_in_* = *d_in_*.*I*, *s_out_* = *d_out_*.*I* are the input and output divisive factors.
- *d_in_ and d_out_* are parameters that control the strength of input and output division and *I* is the output (firing rate) of the inhibitory neuron sending the inhibition.

The value of these parameters chosen were *d_in_* = 0 *and d_out_* = 0.25 consistent with previously published methods (see Fig. 5D in ^44^).

#### Feedback inhibition

In models in which the inhibitory neurons inhibit each other (Fig. 2A, middle column-top; green inhibitory connections), the feedback inhibition was modeled as below using previously published methods^44^.

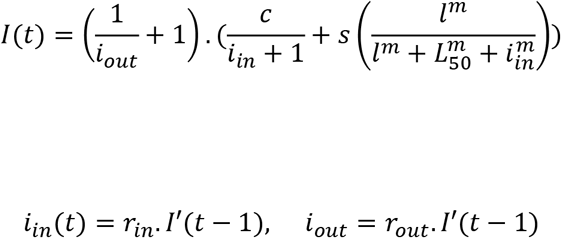

where *I*′ is the output (firing rate) of the other inhibitory neuron at time (*t* − 1).

These equations were iteratively applied until there was no further change in the output of the neurons (i.e., steady state was reached). The time course of reverberant activity due to this feedback connectivity has been plotted in our previously published work^44^.

The values of the feedback parameters used were *r_in_* = 0.8, *r_out_* = 0.01 consistent with previously published methods. We also varied these two parameters (varying feedback; Fig. S1D) to study their effect on categorization index.

#### “Self” inhibition and Donut

In models in which the excitatory neuron receives inhibition from more than one inhibitory neuron (e.g., Fig. 2A, left column top model; baseline model), the inhibition from these sources was combined as below.

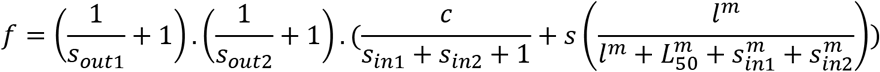

where, 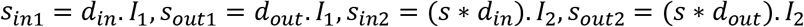

*I*_1_ is the output (firing rate) of the inhibitory neuron 1 and *I*_2_ is the output (firing rate) of the inhibitory neuron 2.

The parameter value *s* controls the strength of self-inhibition, and ranges between 0 and 1. In models that have ‘donut-like connectivity’, *s* is set to 0 (e.g., Fig. 2A, middle column-second from top model; absence of purple projections from Imc neurons to aligned OTid neurons; Donut model). In models which do not have the donut-like connectivity (e.g., Fig. 2A, left column top model; baseline model), *s* is set to 1.

We also vary the value of *s* systematically between 0 (donut) and 1 (maximum self-inhibition) to study the effect of the strength of “self” inhibition on categorization index (Fig. S1I).

#### Recurrent excitation

In the models with recurrent excitation, (of the kind in Fig. S1F), the output of the neuron is scaled by a factor (k; k= 2.5 in Fig. S1GH).

In the model in Fig. 2, the output of the neuron which receives recurrent amplification is modeled as below.

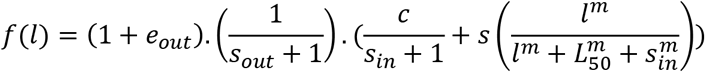

where, *e_out_* = *e_a_*·*A*, *e_a_*= 0.01, and *A* is the output of the neuron sending the amplification.

The value of the parameter is chosen as 0.01 to yield results consistent with the amplification effect of Ipc on OT responses as reported in previously published work (Ipc inactivation results in a 31% decrease in the OTid responses on average^25^). We also vary the amplification factor *e*_*a*_ to study its effect on categorization index (Fig. S1E).

### Models for Figs. 6 and 7

To implement the models in Figs. 6 and 7 (normalization model and donut-like motifs purely via a recurrent route), we used the input-output functions and the effect of divisive inhibition described above. The primary feature of these models that is different from models in Figs. 2, and S1 is that the excitatory neurons send inputs to inhibitory neurons, which then inhibit those excitatory neurons. The output of an excitatory neuron at time t is calculated by applying divisive inhibition (from the inhibitory neurons at time t-1) to its activity at time t-1. This activity is then used to calculate the activity of the inhibitory neurons at time t as below.

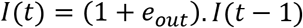

where, *e_out_ e_a_*·*A*, *e_a_* = 0.01, and *A* is the output of the neuron sending the input at time t-1.

This process is repeated iteratively until steady state activity is obtained. The initial activities (at t=0) of the inhibitory neurons are set to their baseline level, and of the excitatory neurons are calculated from the stimulus inputs without any divisive inhibition.

For the models with recurrent amplification (Fig. 7AB), the output of the neuron is scaled by a factor (k, k= 2.5).

### Calculating boundary discriminability and categorization index for the models (Figs. 2, 6, 7, S1 and S4)

For each of the models in Fig 2, 6, 7, S1 and S4 we used a two-stimulus strength morphing protocol described in previously published work^8^. We presented stimulus S1 at location 1 and stimulus S2 at location 2. As the strength of the first stimulus was increased, the strength of the second stimulus was decreased (Figs. 2B, 5E). Responses of the model output neuron (#1) were simulated using this protocol. Random noise from a standard normal distribution was added to the responses. The variance of the noise added depended on the mean value of the responses (m) and a fano factor value (ff): variance = ff *m. This was repeated 30 times (to mimic 30 reps of data collection from a ‘neuron’) and was used to compute the response profiles (mean +/- s.e.m) of that neuron from different model circuits. Similarly, response profiles for 50 neurons were obtained for each model circuit. For all the model runs reported, we used a fano factor value of 6. We also varied the fano factor value to test the effect of noise on model performance (Fig. S1J).

This stimulus presentation protocol results in 2 categories (Category 1: S1 > S2 and Category 2: S1 < S2). We measured the boundary discriminability (bd’) of the responses of neuron 1 by calculating the d-prime between the responses of neuron to stimuli pair straddling either side of the selection boundary and at a distance of 3 units from the boundary as:

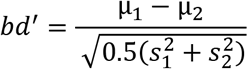

where, μ_1_ and s_1_are the mean and the standard deviation of the responses of the neuron to the stimuli pair from category 1 near the boundary; and μ_2_ and s_2_are the mean and the standard deviation of the responses of the neuron to the stimuli pair from category 2 near the boundary.

To compute the categorization index, we compared two quantities: (modified from previously published work^6,8^) (a) the mean within-category d-prime (WCD’) between the responses of the neuron to pairs of stimulus-pairs (S1 and S2) that are in the same category, and (b) the mean between-category d-prime (BCD’) between the responses of the neuron to pairs of stimulus-pairs that are in different categories. The pairs are chosen while ensuring that (i) the number of pairs used to calculate both these metrics are the same, and (ii) the distribution of the distances between the chosen pairs for calculating both the metrics are the same. The categorization index is calculated from these two metrics as:

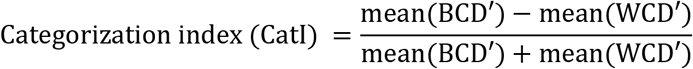

CatI= 1 indicates idealized, step-like responses; = 0 indicates linear, non-categorical responses; < 0 indicates better discriminability within category than between categories (Fig. S1C).

### Data collection protocol

Visual stimuli used here have been described previously ^8,49^. Briefly, looming visual dots are flashed at different locations on a tangent TV monitor in front of the owl. Looming stimuli were dots that expanded linearly in size over time, starting from a size of 0.6° in radius. The speed of the loom was decided based on the stimulus protocol and varied between 9.6°/s and 21.6°/s. Visual stimuli were presented for a duration of 250ms with an inter stimulus interval of 1.25s to 4s.

#### Receptive fields (RFs)

For measuring spatial RFs of Imc, OTid and Ipc neurons, a single stimulus of a fixed contrast was presented at the sampled locations. The order of locations at which the stimulus was presented was randomized to minimize adaptation. A location is considered to be inside the RF if it evokes responses significantly different from that of baseline response. All other locations are considered to be outside the RF.

The following stimulus protocols were used for measuring data reported in Figs. 3, 4.

## I. Paired OTid and Imc data collection

### 1. “Other” inhibition

To measure the strength of “other” inhibition sent from an Imc neuron to a distant location in the OTid space map, we simultaneously recorded from the Imc neuron (with a multi-barrel glass electrode filled with the kynurenic acid), and a spatially misaligned “other” OTid neuron (with a tungsten electrode) as described below; we ensured that the half-max of the RF of the OTid neuron lay outside (did not overlap with) the half-max of the Imc RF.

For measuring “other” inhibition, we recorded the following data curves together in an interleaved manner.

a. Tuning curve (TC) centered at the OTid RF peak: A 1-dimensional spatial tuning curve centered at the peak of the OTid RF. The stimulus had a loom strength of 9.6°/s.
b. Tuning curve centered at the OTid RF peak + competing stimulus centered at the Imc RF peak (TCC): The same curve as in a), but along with a (distant) competitor positioned at the peak of the Imc RF. The strength of the competing stimulus was chosen to be 19.2°/s. This second competing stimulus drives the Imc neuron, which sends strong competitive inhibition to the OTid neuron. Since the competing stimulus was more salient than the stimulus driving the OTid neuron, the responses of the OTid neuron were strongly suppressed consistent with previous published results^49^.
c. Tuning curve (TC) centered at the Imc RF peak: 1-dimensional spatial tuning curve centered at the Imc RF. The loom strength of the stimulus was chosen to be 9.6°/s.

We measured the above 3 curves both when the Imc neuron is intact (‘baseline’ condition) and focally inactivated (‘inactivation’ condition) and compare the responses as below. In a subset of the data, we also measured the curves after the responses of the Imc neurons recovered (Fig. S2A, ‘recovery’ condition) from focal iontophoretic inactivation. Inactivation curves were measured 15 min after the drug ejection was started. Recovery curves were measured 15 min after the drug ejection was stopped.

We analyzed the data from these three curves as below.

First, a distant competing stimulus is known to typically suppress OTid responses^49,50^, something that we confirmed by comparing the responses of OTid neurons to the TC (curve a) and TCC (curve b); Fig. S2B. Notably, it is also known that some OTid neurons do not show such suppression, and any such neurons were excluded from our analyses^9^.

Next, to measure the amount of Imc inactivation, we compared the responses of the Imc neurons to the TC measured at the Imc RF peak (curve c) in the baseline condition versus the inactivation condition. Kynurenic acid was able to effectively shut down Imc responses: (Fig. S2C; red, median strength of Imc inactivation = 95%, 95% CI of median = [87%, 103%], p = 3.8e-6, sign test, n=19).

Finally, to measure the strength of “other” inhibition, we compared the TCC responses (curve b) in the baseline condition and the inactivation condition. In the baseline condition, the Imc neuron was intact, driven by the competing stimulus and sent strong inhibition to the OTid neuron. In the inactivation condition, the Imc neuron was silenced, as a result of which the OTid neuron was released from inhibition and exhibited an increase in responses.

Any observed increase in OTid responses quantified the amount of suppression due to Imc onto the OTid location, thereby directly estimating the strength of net “other” inhibition: % change in OTid responses = 100* (responses in Imc intact condition – responses in Imc off condition)/responses in Imc off condition. To obtain an accurate estimate of the % change, we considered the responses to stimulus S1 (in the two conditions) at all locations inside the RF as follows. We fit a straight line to the plot of OTid responses to S1 in the Imc intact vs. Imc of conditions (Fig. 3F), calculated the slope of the best-fit line, and used it to compute the average value of % change as: %change in OTid responses = 100 * (slope-1). This procedure was also used to quantify “self” inhibition, “gap” inhibition and “different lobe” inhibition (below).

### 2. “Self” inhibition

To measure the strength of “self” inhibition sent from an Imc neuron to a matched location in the OTid space map, we simultaneously recorded from the Imc neuron (with a multi-barrel glass electrode filled with the kynurenic acid), and a spatially aligned “self” OTid neuron (with a tungsten electrode) as described below; we ensured that the half-max of the RF of the OTid neuron overlapped with the half-max of the Imc RF.

We recorded a spatial tuning curve centered at the peak of the OTid (and therefore, Imc) RF. The loom speed of the stimulus used was 9.6°/s; this stimulus drives both the OTid and the Imc neuron.

We compared the responses of the OTid neuron in the Imc-intact (baseline) and Imc inactivated condition and the difference quantified the strength of “self” inhibition (Fig. 3L, blue).

To measure the amount of Imc inactivation, we compared the responses of the Imc neuron in the baseline and the inactivation condition as before. Kynurenic acid was able to effectively shut down Imc responses: (median strength of Imc inactivation = 92%, 95% CI of median = [86% 98%], p = 7.5e-9, sign test).

### 3. “Gap” inhibition

To measure the strength of “self” inhibition sent from an Imc neuron with a multilobed RF to the locations in the OTid space map between the RF lobes (gaps), we simultaneously recorded from the Imc neuron (with a multi-barrel glass electrode filled with the kynurenic acid), and “gap” OTid neuron (with a tungsten electrode), the RF (half-max) of which was located in the gap between the half-max of the lobes of the Imc RF (Fig. 4A-C). We recorded the same set of curves as in the “other” inhibition case (see above) from the gap OTid neuron, and performed similar analyses on the data to quantify the strength of “gap” inhibition.

### 4. “Different lobe” inhibition

To measure the strength of inhibition sent “from” locations within one lobe of an Imc neuron with a multilobed RF to the locations in the OTid space map within other lobes of the Imc RF (“different lobe” inhibition), we simultaneously recorded from a multilobe Imc neuron (with a multi-barrel glass electrode filled with the kynurenic acid), and an OTid neuron (with a tungsten electrode), the RF (half-max) of which overlapped with one of the lobes of the Imc RF (Fig. 4G-I). Then we measured the same set of curves as in the “other” case while ensuring one additional detail. We recorded the TC curve (curve a) with stimulus (S1) centered at the OTid RF peak (and therefore within one of the lobes of the Imc RF as well). For the TCC (curve b), the second stimulus S2 was centered at the peak of a different lobe of the Imc neuron’s multilobe RF. Thus, both stimuli excited the Imc neuron (during curve b) as they lay within its RF, but only S1 excited the OTid neuron and S2 served as a (distant) competitor from the its perspective.

An OTid neuron in this configuration was defined as a “different lobe” OTid neuron; the half max of its RF overlapped with the half max of one of the lobes of the Imc RF, but not of other lobes. (Note that by definition, every “different lobe” OTid neuron is also a potentially “self” OTid neuron, and admits to the use of the appropriate stimulus protocol to measure “self” inhibition. However, the opposite is not true, since “self” OTid neurons can be identified for Imc neurons with single lobed RFs as well, but “different lobe” OTid neurons are only defined for multilobe Imc neurons).

We applied similar analyses as in the “other” case to the data from “different lobe” OTid neurons and quantified the strength of “different lobe” inhibition.

## II. Paired OTid and Ipc data collection (Figs. 5, S4)

To test if the donut-like inhibitory motif is required for (robustness-to-noise and) categorization, we made paired recordings at spatially aligned Ipc and OTid neurons. We used a strength morphing stimulus protocol described in previously published work^8^. Briefly, we presented one stimulus (S1) inside the RF of the OTid neuron (and also Ipc neuron because their RFs overlap). Simultaneously, we presented a competing stimulus (S2) 30° away along azimuth from S1. As the strength of the S1 decreased, the strength of S2 stimulus increased (Fig. 5E).

We applied the same analyses described above for the model in Figs. 2, S1 to compute the categorization index for experimental data reported in Fig. 5.

### Data analyses and statistical tests

All analyses were carried out with custom MATLAB code. Parametric or non-parametric statistical tests were applied based on whether the distributions being compared were Gaussian or not, respectively (Lilliefors test of normality). The Holm-Bonferroni correction was used to account for multiple comparisons; the abbreviation “with HBMC correction” in the text stands for “with Holm-Bonferroni correction for multiple comparisons”. All tests were two-sided. Data shown as a ± b refer to mean ± standard deviation, unless specified otherwise. The ‘*’ symbol indicates significance at the 0.05 level (after corrections for multiple comparisons, if applicable). Correlations were tested using Pearson’s correlation coefficient (*corr* command in MATLAB with the ‘Pearson’ option).

### Code and data availability

Software code and the data that support the findings of this study are available from the corresponding author upon reasonable request.

## SUPPLEMENTAL FIGURES (S1-S4)

**Fig. S1.**
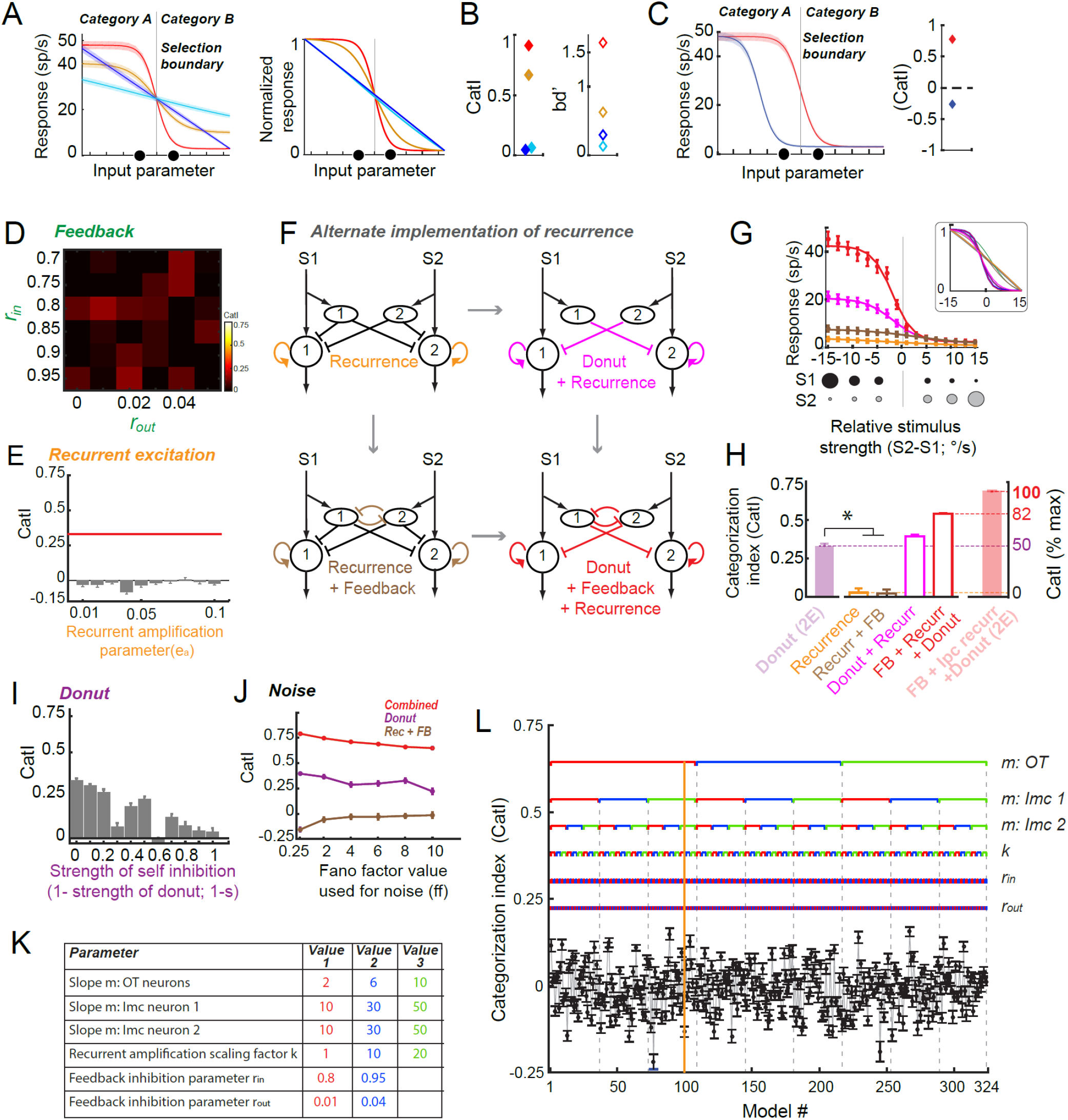
Donut-like inhibition surpasses other circuit motifs in its ability to generate categorical representations over a range of values of key model parameters and amounts of response noise (modeling). **(A)** <u>Left:</u> Schematic showing 4 different mathematically generated response profiles, as a function of continuously varying input; conventions as in Fig. 1A. Red and orange profiles are both categorical; orange transitions less abruptly than red, and additionally, orange is a scaled down version of red. Blue and cyan profiles are both linear (equally non-categorical); cyan is a scaled down version of blue. <u>Right</u>: Response profiles in left panel, normalized between 0 and 1; only means are shown. Red profile is clearly seen to transition more abruptly than orange, blue and cyan profiles are clearly seen to transition equally gradually (in a linear manner). **(B)** <u>Left</u> (filled symbols): Categorization index (CatI) for response profiles in A (Methods). It is sensitive to the abruptness of the transition in response profiles – it is greater for red than orange, and they are both greater than blue. Notably, CatI is insensitive to scaling – it is nearly equal for blue and cyan. <u>Right</u>: By contrast, discriminability computed across the selection boundary, bd’, is confounded by scaling – it is greater for blue than cyan (and the difference between its values for red and orange is greater than for CatI, due to the scaling factor between red and orange profiles in A, left). It is therefore not a reliable metric of categorization. **(C)** <u>Left:</u> Schematic comparing two mathematically generated sigmoidal response profiles as a function of a continuously varying input; conventions as in Fig 1A. Red – response profile in which the response transition from high to low levels occurs at the selection (or category) boundary, over a small range of input values. Dark blue – a similarly abrupt response profile in which the response transition occurs far from the selection boundary. Gray vertical line: ideal selection (or category) boundary. Translucent band: variability in responses; fano factor of 6 used to generate these responses (Methods). <u>Right:</u> Categorization index (CatI) characterizes strength of categorization of response profiles in left panel. CatI value is positive and high for response profiles (red) that exhibit an abrupt transition across the ideal boundary. However, it is negative for response profiles (blue) that exhibit a transition far from the boundary (even when the transition is abrupt), because the within category discriminability (WCD) is higher than the between category discriminability (BCD)). **(D)** Effect of varying strength of feedback inhibition in model in Fig. 2A middle-column top, on CatI. rin: input divisive factor; rout: output divisive factor (Methods). Range of variation based on previously published work^44^. Maximum CatI = 0.157 (lower than from the donut-like motif, CatI = 0.331). **(E)** Effect of varying strength of recurrent amplification in model in Fig. 2A, left column-bottom on CatI. Red line: value from circuit with donut-like motif only. CatI does not change systematically (corr= −0.14; p =0.54; corr test), indicating that varying the strength of amplification does not affect the CatI estimate for this motif in Fig. 2D. **(F-H)** Computational models with alternate implementation of Ipc-recurrent amplification, and termed ‘recurrent amplification’ (similar to that employed in the modeling literature). Here, unlike Ipc-recurrent amplification in Fig. 2A, recurrent amplification within each channel is not under the control of powerful competitive inhibition. Nonetheless, just like in Fig. 2D, recurrent amplification by itself, or in conjunction with feedback, is ineffective at producing categorization (i.e., CatI values are low). (F) Model circuits with recurrence alone (top-left), recurrence and donut (top-right), recurrence and feedback (bottom-left), and all three motifs (bottom-right). (G) Plots of the response profiles to strength-morphing protocol (and normalized response profiles; inset), obtained from the four model circuits in F. (H) Plots of CatI of these response profiles. All conventions as in Fig. 2CD. **(I)** Effect of varying strength of ‘self’-inhibition in model in Fig. 2A, left column-top on CatI. Corr = −0.81, p = 3e-3 (correlation test). Maximal effect on CatI is when ‘self’-inhibition=0, i.e., when the circuit has donut-like inhibition. **(J)** Comparison (across three key models) of CatI from simulated response profiles as a function of fano-factor. Models are: circuit with donut-like inhibitory motif only (Fig. 2A, middle column-second from top), circuit with Ipc-recurrent amplification and feedback inhibition together (Fig. 2A, middle column-second from bottom), and circuit with all three motifs combined (Fig. 2A, right column-bottom). **(K, L)** Exploration of the efficacy of models without the donut-like motif (but with feedback inhibition and recurrent amplification) in achieving categorical responses, when values of various key model parameters are varied. The model architecture used in these simulations is the same as that in F, bottom-left. (K) Table showing parameters that are varied and their values; color codes signify different values of each parameter. A total of 324 combinations of the parameter values are explored. (L) CatI of response profiles obtained from each of these 324 circuit models; mean ± s.e.m, n=50 model neurons. Each point in the plot corresponds to a combination of parameter values. The brackets above the CatI plot indicate the color-coded value that each parameter takes for that model; the color codes are consistent with the table in (K); vertical dashed lines have been added to aid visualization. Vertical orange line highlights model #100; for this model, m (OT) = 2 (red), m (Imc1) =50 (green), m (Imc2) = 50 (green); k = 1 (red); rin =0.95 (blue); rout =0.04 (blue).

**Fig. S2.**
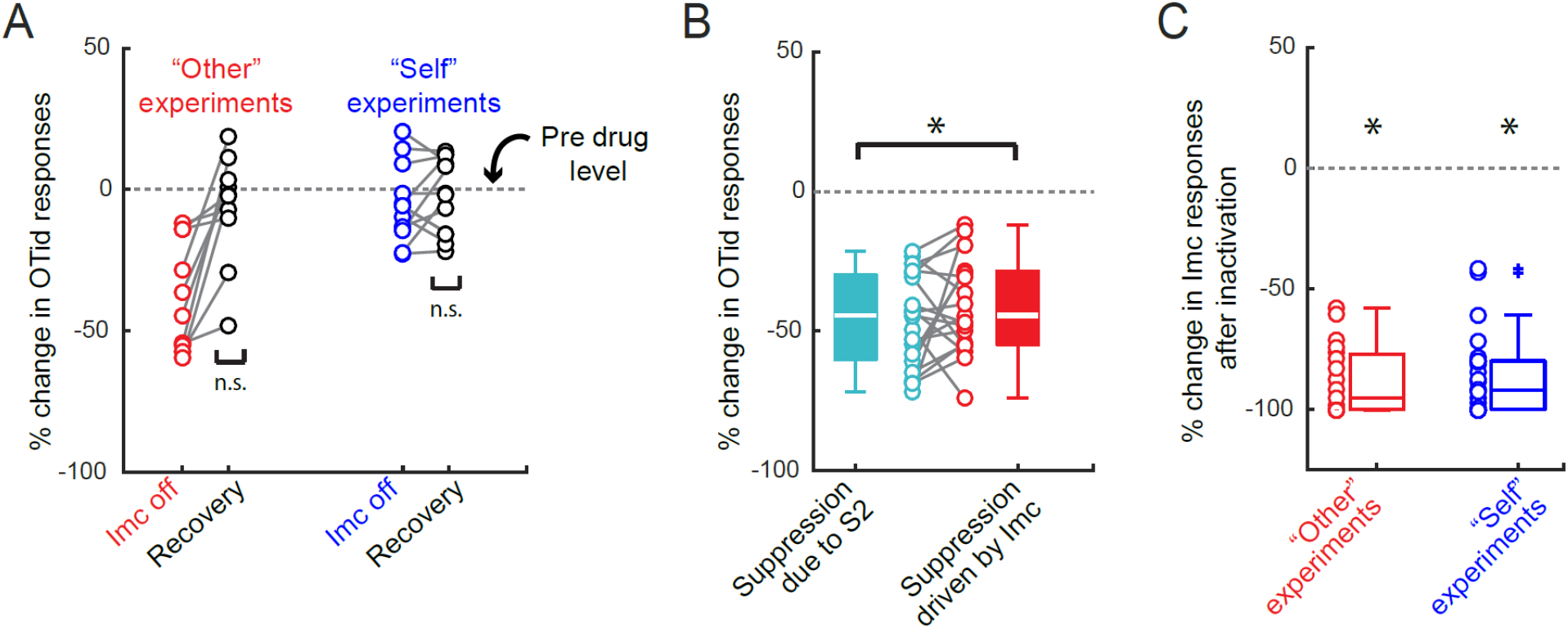
Supporting experimental data for Figure 3. **(A)** Recovery of OTid responses from kynurenic acid iontophoresis for experiments in Fig. 3. <u>Data on left</u>: “Other” experiments. OTid responses revealed significant change during iontophoresis (red dots; subset of the data reproduced from Fig. 3L), but returned to pre-drug baseline (horizontal line) in recovery (black dots; p=0.32, t-test against 0). Recovery data obtained 15 min after iontophoretic eject current was switched to retain current; Methods). Data show recovery, demonstrating that the effects reported Fig. 3 are due specifically to drug iontophoresis/Imc inactivation. <u>Data on right</u>: “Self” experiments. OTid responses showed no significant change during iontophoresis (blue dots; reproduced from Fig. 3L), and stayed around zero in recovery (black dots; p=0.58, t-test against 0. **(B)** “Other” experiment. Comparison of suppression provided by Imc with that due to S2 (i.e., the maximum amount of suppression experienced by the OTid neuron in this stimulus protocol). This is done by comparing (i) % suppression of OTid responses by stimulus S2 when Imc is intact, computed by comparing OTid responses to S1 alone versus the paired presentation of S1 and S2 (teal^49^), with (ii) % suppression of OTid responses to paired presentation of S1 and S2 produced by Imc inactivation, computed by comparing OTid responses in the Imc-intact to Imc-off conditions (red; data reproduced from Fig. 3L). Consistent with^32^, nearly all the suppression due to competitor S2 (teal) is supplied by Imc (red) (p= 0.22, t-test, teal vs. red), verifying that Imc supplies powerful “other” inhibition. **(C)** Quantifying the effectiveness of Imc inactivation by kynurenic acid iontophoresis in the “other” (red) and “self”-inhibition (blue) experiments. In both cases, the inactivation was highly effective. Red: median % change in response; median = 95%, 95% CI of median = [87%, 103%], p = 3.8 e-6, sign test; Blue: median % change in responses median =0.92, 95% CI of median = [86%, 98%], p = 7.45e-9, sign test.

**Fig. S3.**
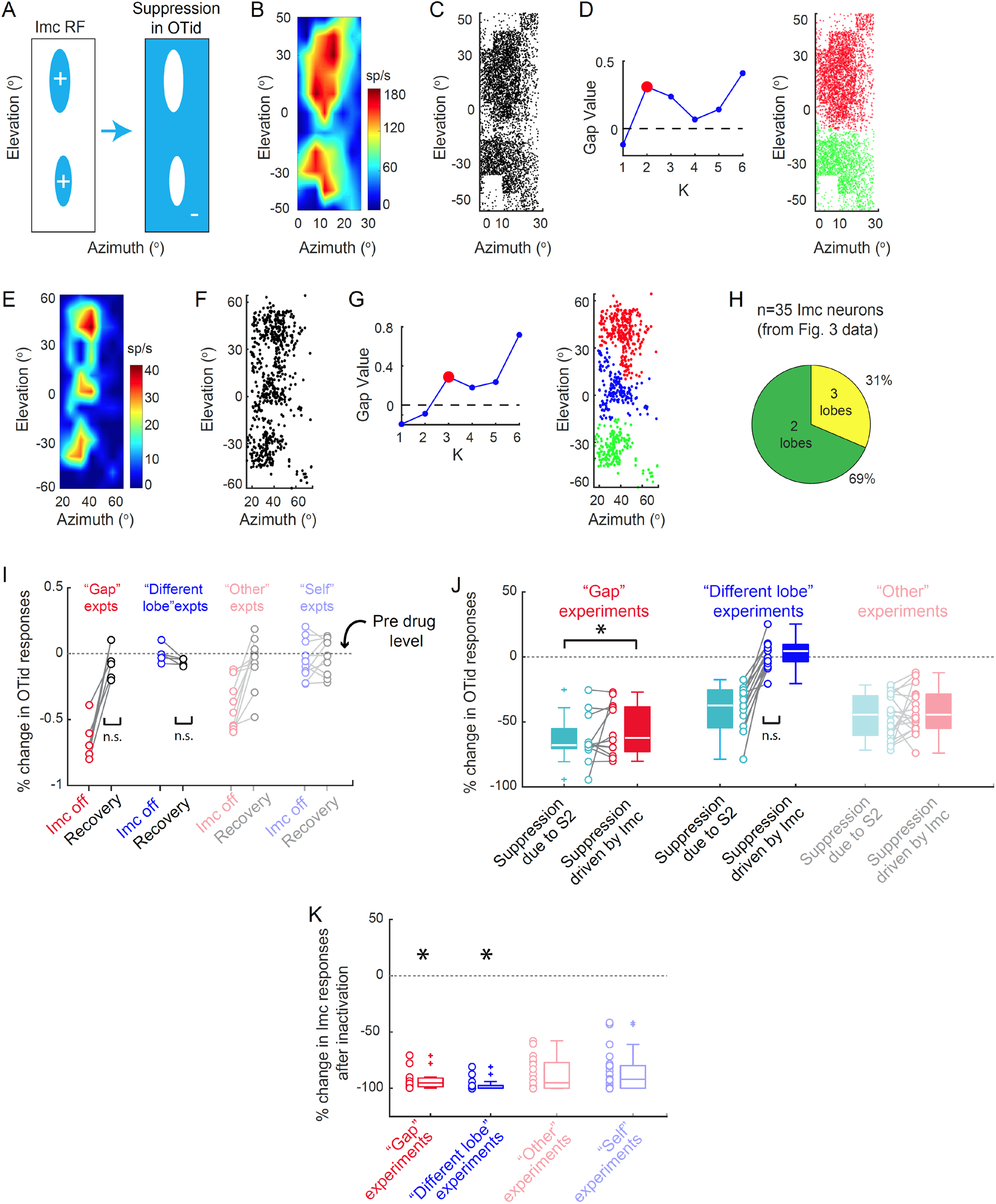
Detecting the number of Imc RF lobes, and other supporting experimental data for Figure 4. no inhibition arrives due to the neuron on the left. **(B)** Imc RF reproduced from Fig. 4A-left. **(C)** Firing rate map from B converted to density of points in 2-D plane following published procedures^52^. **(D)** <u>Left</u>: Density peaks clustering method^89^ is applied to the data in B, forcing the method to yield either 1 cluster, 2 clusters, … up to 6 clusters. Following that, the gap statistic model selection metric^90^ is applied to the clustering results to identify the optimal number of clusters in the data. The optimal number is the number for which the “gap value” exceeds 0 for the first time; here it is 2. <u>Right</u>: Same as B, but with the two colors indicating the two distinct lobes identified by the clustering method (+ gap statistic) as the optimal two best clusters in the RF data. **(E-G)** Same as B-D, but for Imc RF in Fig. 4G-left; determined to be three-lobed. **(H)** Pie-chart showing proportion of two-lobed vs. three-lobed RFs across all the multi-lobed Imc neurons recorded in these experiments (Fig. 4). **(I)** Recovery of OTid responses from kynurenic acid iontophoresis for experiments in Fig. 4. Conventions as in Fig. S2A; data from Fig. S2A reproduced here for comparison. Data show recovery (“Gap” experiment: p =0.39, t-test of black data against 0; “Other” lobe experiment: p=0.29, ranksum test of black data against 0), demonstrating that any effects reported Fig. 4 are due specifically to drug iontophoresis/Imc inactivation. **(J)** Comparison of suppression provided by Imc with that due to S2 (i.e., the maximum amount of suppression experienced by the OTid neuron in this stimulus protocol) in the “gap” (red data) and “different lobe” (blue data) experiments. Conventions as in Fig. S2B; data from Fig. S2B reproduced here for comparison. “<u>Gap” experiment</u>: Consistent with^32^, nearly all the suppression due to competitor S2 (teal) is supplied by Imc (red) (p= 0.47, teal vs. red, ranksum test). <u>“Different lobe” experiment</u>: The Imc neuron that is being inactivated does not provide any of the inhibition to the OTid neuron due to S2 (p = 0.35, blue dots against 0, t-test). This clearly demonstrates, consistent with the predictions of ^52^, that a different Imc neuron exists, which has one RF lobe encoding S2’s location, but other RF lobes (if there are others for that neuron) not encoding S1’s location, thereby delivering inhibition to that location. This is one of the ‘signature’ properties for the combinatorial encoding of space described in that study. **(K)** Quantifying the effectiveness of Imc inactivation by kynurenic acid iontophoresis in the “gap” (red) and “different lobe” (blue) experiments. “Gap” experiment: p =1.13 e-12 t-test against 0; “Other” lobe experiment: p =1.53e-5, signtest. Conventions as in Fig. S2C; data from Fig. S2C reproduced here for comparison.

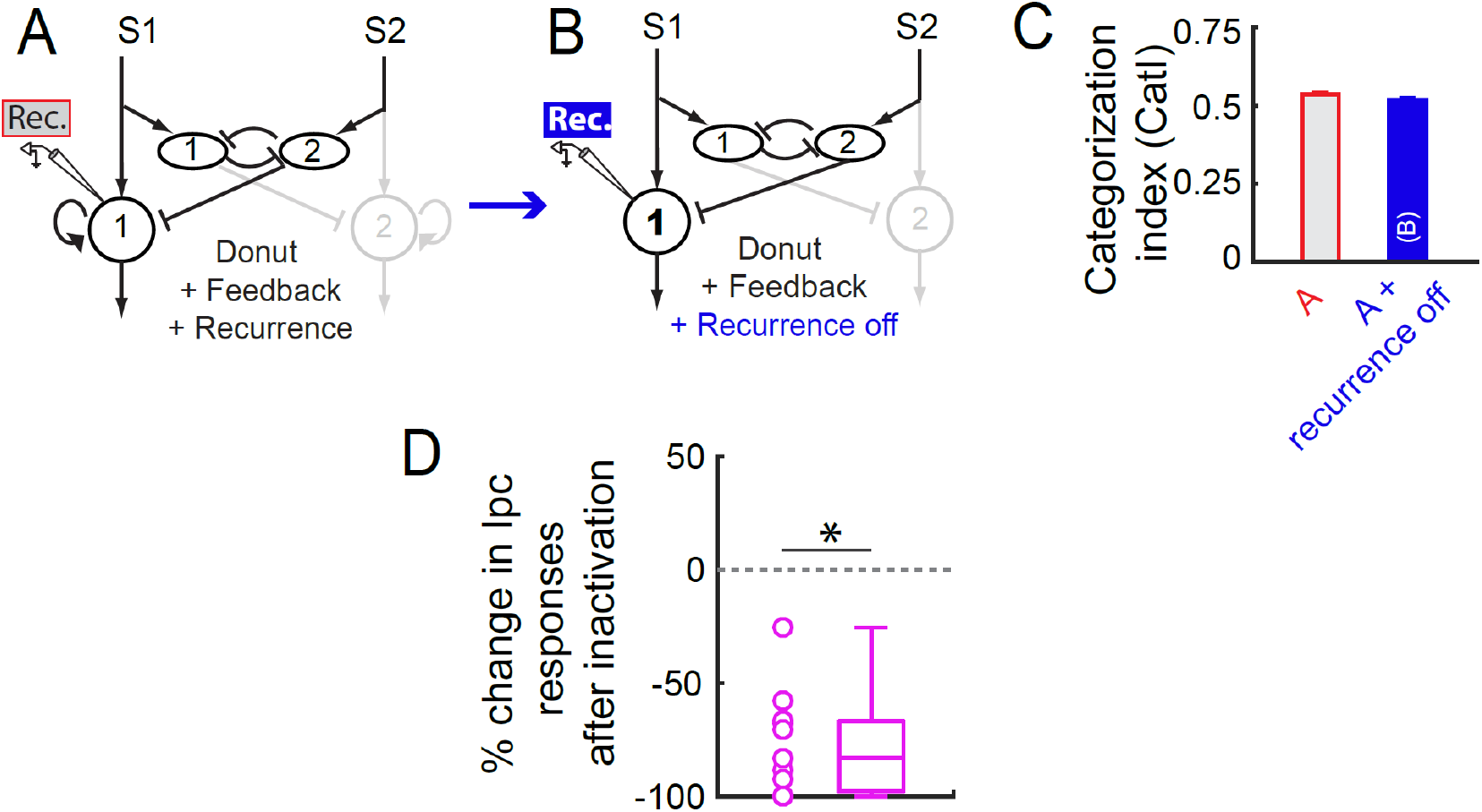
(A-C) Modeling: Effect of silencing recurrent amplification. **(A)** Computational model of modified version of the midbrain selection network; reproduced from Fig. S1F, bottom-right. Recall that this network has donut-like inhibition, feedback inhibition, and recurrent amplification (just like the midbrain selection network), but with the one modification that the recurrent amplification (black curved arrow) is not under the control of competitive inhibition. The strength of curved black arrow is not affected by inhibition from oval neurons, in contrast to ‘Ipc-recurrent’ amplification in the midbrain selection network, which is (Fig. 5A). Incidentally, this form of recurrence, termed simply, ‘recurrent’ amplification, represents the typical implementation in published models of selection. **(B)** Model in A, but with recurrent amplification silenced. (C) Plots of CatI of the response profiles obtained from model circuits in A (red-gray) and B (blue) presented with the strength-morphing protocol. These results show that silencing just recurrent amplification (when it is not under the control of competitive inhibition, B), causes no discernible impact on categorization (C, blue vs. red-gray). (This is in contrast to inactivation of Ipc in the barn owl midbrain network, which causes abolishment of categorization, just as introducing self-inhibition or ‘filling- in’ the donut-hole does; Fig. 5D, pink vs. gold. Thus, the proposed focal Ipc inactivation in the barn owl midbrain network (Fig. 5C) is akin to introducing self-inhibition onto Ipc (Fig. 5B) rather than to simply silencing recurrent amplification.) All conventions as in Fig. 5. **(D) Experiments: Ipc is effectively inactivated in experiments in Fig. 5.** Mean suppression of Ipc responses upon iontophoresis of kynurenic acid was 77.42% with a standard deviation of 22.82%; p-value = 5.3e-7; t-test against 0.

## REFERENCES

1 Freedman, D. J. & Assad, J. A. A proposed common neural mechanism for categorization and perceptual decisions. Nat Neurosci 14, 143–146 (2011).

2 Niessing, J. & Friedrich, R. W. Olfactory pattern classification by discrete neuronal network states. Nature 465, 47–52 (2010).

3 Leopold, D. A. & Logothetis, N. K. Multistable phenomena: changing views in perception. Trends Cogn Sci 3, 254–264, doi:S1364-6613(99)01332-7 [pii] (1999).

4 Gollisch, T. & Meister, M. Eye smarter than scientists believed: neural computations in circuits of the retina. Neuron 65, 150–164 (2010).

5 Jiang, X., Chevillet, M. A., Rauschecker, J. P. & Riesenhuber, M. Training Humans to Categorize Monkey Calls: Auditory Feature- and Category-Selective Neural Tuning Changes. Neuron 98, 405–416 e404, doi:10.1016/j.neuron.2018.03.014 (2018).

6 Freedman, D. J. & Assad, J. A. Experience-dependent representation of visual categories in parietal cortex. Nature 443, 85–88 (2006).

7 Freedman, D. J., Riesenhuber, M., Poggio, T. & Miller, E. K. Categorical representation of visual stimuli in the primate prefrontal cortex. Science 291, 312–316, doi:10.1126/science.291.5502.312 (2001).

8 Mysore, S. P. & Knudsen, E. I. Flexible categorization of relative stimulus strength by the optic tectum. J Neurosci 31, 7745–7752 (2011).

9 Mysore, S. P., Asadollahi, A. & Knudsen, E. I. Signaling of the strongest stimulus in the owl optic tectum. J Neurosci 31, 5186–5196 (2011).

10 Hirokawa, J., Vaughan, A., Masset, P., Ott, T. & Kepecs, A. Frontal cortex neuron types categorically encode single decision variables. Nature 576, 446–451, doi:10.1038/s41586-019-1816-9 (2019).

11 Bathellier, B., Ushakova, L. & Rumpel, S. Discrete neocortical dynamics predict behavioral categorization of sounds. Neuron 76, 435–449, doi:10.1016/j.neuron.2012.07.008 (2012).

12 Xin, Y. et al. Sensory-to-Category Transformation via Dynamic Reorganization of Ensemble Structures in Mouse Auditory Cortex. Neuron 103, 909–921 e906, doi:10.1016/j.neuron.2019.06.004 (2019).

13 Jovanic, T. et al. Competitive Disinhibition Mediates Behavioral Choice and Sequences in Drosophila. Cell 167, 858–870 e819, doi:10.1016/j.cell.2016.09.009 (2016).

14 Mysore, S. P. & Knudsen, E. I. The role of a midbrain network in competitive stimulus selection. Curr Opin Neurobiol 21, 653–660, doi:10.1016/j.conb.2011.05.024 (2011).

15 Carello, C. D. & Krauzlis, R. J. Manipulating intent: evidence for a causal role of the superior colliculus in target selection. Neuron 43, 575–583 (2004).

16 Cavanaugh, J. & Wurtz, R. H. Subcortical modulation of attention counters change blindness. J Neurosci 24, 11236–11243 (2004).

17 Lovejoy, L. P. & Krauzlis, R. J. Inactivation of primate superior colliculus impairs covert selection of signals for perceptual judgments. Nat Neurosci 13, 261–266 (2010).

18 McPeek, R. M. & Keller, E. L. Deficits in saccade target selection after inactivation of superior colliculus. Nat Neurosci 7, 757–763 (2004).

19 Muller, J. R., Philiastides, M. G. & Newsome, W. T. Microstimulation of the superior colliculus focuses attention without moving the eyes. Proc Natl Acad Sci U S A 102, 524–529 (2005).

20 Krauzlis, R. J., Lovejoy, L. P. & Zenon, A. Superior colliculus and visual spatial attention. Annu Rev Neurosci 36, 165–182 (2013).

21 Knudsen, E. I. Control from below: the role of a midbrain network in spatial attention. Eur J Neurosci 33, 1961–1972 (2011).

22 Mysore, S. P. & Knudsen, E. I. Descending control of neural bias and selectivity in a spatial attention network: rules and mechanisms. Neuron 84, 214–226 (2014).

23 Fecteau, J. H. & Munoz, D. P. Salience, relevance, and firing: a priority map for target selection. Trends Cogn Sci 10, 382–390 (2006).

24 Nummela, S. U. & Krauzlis, R. J. Inactivation of primate superior colliculus biases target choice for smooth pursuit, saccades, and button press responses. J Neurophysiol 104, 1538–1548 (2010).

25 Asadollahi, A. & Knudsen, E. I. Spatially precise visual gain control mediated by a cholinergic circuit in the midbrain attention network. Nature communications 7, 13472, doi:10.1038/ncomms13472 (2016).

26 Wang, Y., Luksch, H., Brecha, N. C. & Karten, H. J. Columnar projections from the cholinergic nucleus isthmi to the optic tectum in chicks (Gallus gallus): a possible substrate for synchronizing tectal channels. J Comp Neurol 494, 7–35 (2006).

27 Marin, G., Mpodozis, J., Sentis, E., Ossandon, T. & Letelier, J. C. Oscillatory bursts in the optic tectum of birds represent re-entrant signals from the nucleus isthmi pars parvocellularis. J Neurosci 25, 7081–7089 (2005).

28 Schryver, H. M., Straka, M. & Mysore, S. P. Categorical Signaling of the Strongest Stimulus by an Inhibitory Midbrain Nucleus. J Neurosci 40, 4172–4184, doi:10.1523/JNEUROSCI.0042-20.2020 (2020).

29 Schryver, H. M., Lim, J.-X. & Mysore, S. P. Distinct neural mechanisms construct classical versus extraclassical inhibitory surrounds in an inhibitory nucleus in the midbrain attention network. (in revision, Nat Commun), bioRxiv doi:https://doi.org/10.1101/2020.03.13.990952 (2020).

30 Schryver, H. M. & Mysore, S. P. Spatial Dependence of Stimulus Competition in the Avian Nucleus Isthmi Pars Magnocellularis. Brain Behav Evol 93, 137–151, doi:10.1159/000500192 (2019).

31 Wang, Y., Major, D. E. & Karten, H. J. Morphology and connections of nucleus isthmi pars magnocellularis in chicks (Gallus gallus). J Comp Neurol 469, 275–297 (2004).

32 Mysore, S. P. & Knudsen, E. I. A shared inhibitory circuit for both exogenous and endogenous control of stimulus selection. Nat Neurosci 16, 473–478, doi:10.1038/nn.3352 (2013).

33 Marin, G. et al. A cholinergic gating mechanism controlled by competitive interactions in the optic tectum of the pigeon. J Neurosci 27, 8112–8121 (2007).

34 Mysore, S. P. & Kothari, N. B. Mechanisms of competitive selection: A canonical neural circuit framework. Elife 9, doi:10.7554/eLife.51473 (2020).

35 Olsen, S. R., Bhandawat, V. & Wilson, R. I. Divisive normalization in olfactory population codes. Neuron 66, 287–299 (2010).

36 Bollimunta, A. & Ditterich, J. Local computation of decision-relevant net sensory evidence in parietal cortex. Cereb Cortex 22, 903–917, doi:10.1093/cercor/bhr165 (2012).

37 Hartline, H. K., Wagner, H. G. & Ratliff, F. Inhibition in the eye of Limulus. J Gen Physiol 39, 651–673 (1956).

38 Engel, T. A., Chaisangmongkon, W., Freedman, D. J. & Wang, X. J. Choice-correlated activity fluctuations underlie learning of neuronal category representation. Nature communications 6, 6454, doi:10.1038/ncomms7454 (2015).

39 Wang, X. J. Neural dynamics and circuit mechanisms of decision-making. Curr Opin Neurobiol 22, 1039–1046, doi:10.1016/j.conb.2012.08.006 (2012).

40 Machens, C. K., Romo, R. & Brody, C. D. Flexible control of mutual inhibition: a neural model of two-interval discrimination. Science 307, 1121–1124 (2005).

41 Mysore, S. P. & Kothari, N. B. Mechanisms of competitive selection: A canonical neural circuit framework In review

42 Wang, X. J. Decision making in recurrent neuronal circuits. Neuron 60, 215–234 (2008).

43 Chen, Q., Pei, Z., Koren, D. & Wei, W. Stimulus-dependent recruitment of lateral inhibition underlies retinal direction selectivity. eLife 5, doi:10.7554/eLife.21053 (2016).

44 Mysore, S. P. & Knudsen, E. I. Reciprocal inhibition of inhibition: a circuit motif for flexible categorization in stimulus selection. Neuron 73, 193–205 (2012).

45 Goddard, C. A., Mysore, S. P., Bryant, A. S., Huguenard, J. R. & Knudsen, E. I. Spatially Reciprocal Inhibition of Inhibition within a Stimulus Selection Network in the Avian Midbrain. PLoS One 9, e85865 (2014).

46 Albantakis, L. & Deco, G. The encoding of alternatives in multiple-choice decision making. Proc Natl Acad Sci U S A 106, 10308–10313, doi:10.1073/pnas.0901621106 (2009).

47 Sereno, M. I. & Ulinski, P. S. Caudal topographic nucleus isthmi and the rostral nontopographic nucleus isthmi in the turtle, Pseudemys scripta. J Comp Neurol 261, 319–346 (1987).

48 Asadollahi, A., Mysore, S. P. & Knudsen, E. I. Rules of competitive stimulus selection in a cholinergic isthmic nucleus of the owl midbrain. J Neurosci 31, 6088–6097 (2011).

49 Mysore, S. P., Asadollahi, A. & Knudsen, E. I. Global inhibition and stimulus competition in the owl optic tectum. J Neurosci 30, 1727–1738 (2010).

50 Rizzolatti, G., Camarda, R., Grupp, L. A. & Pisa, M. Inhibitory effect of remote visual stimuli on visual responses of cat superior colliculus: spatial and temporal factors. J Neurophysiol 37, 1262–1275 (1974).

51 Knudsen, E. I. Auditory and visual maps of space in the optic tectum of the owl. J. Neurosci. 2, 1177–1194 (1982).

52 Mahajan, N. R. & Mysore, S. P. Combinatorial Neural Inhibition for Stimulus Selection across Space. Cell reports 25, 1158–1170 e1159, doi:10.1016/j.celrep.2018.10.022 (2018).

53 Fadok, J. P. et al. A competitive inhibitory circuit for selection of active and passive fear responses. Nature 542, 96–100, doi:10.1038/nature21047 (2017).

54 Mante, V., Sussillo, D., Shenoy, K. V. & Newsome, W. T. Context-dependent computation by recurrent dynamics in prefrontal cortex. Nature 503, 78–84 (2013).

55 Chaisangmongkon, W., Swaminathan, S. K., Freedman, D. J. & Wang, X. J. Computing by Robust Transience: How the Fronto-Parietal Network Performs Sequential, Category-Based Decisions. Neuron 93, 1504–1517 e1504, doi:10.1016/j.neuron.2017.03.002 (2017).

56 Carandini, M., Heeger, D. J. & Movshon, J. A. Linearity and normalization in simple cells of the macaque primary visual cortex. J Neurosci 17, 8621–8644 (1997).

57 Kouh, M. & Poggio, T. A canonical neural circuit for cortical nonlinear operations. Neural Comput 20, 1427–1451, doi:10.1162/neco.2008.02-07-466 (2008).

58 Louie, K., Khaw, M. W. & Glimcher, P. W. Normalization is a general neural mechanism for context-dependent decision making. Proc Natl Acad Sci U S A 110, 6139–6144, doi:10.1073/pnas.1217854110 (2013).

59 Busse, L., Wade, A. R. & Carandini, M. Representation of concurrent stimuli by population activity in visual cortex. Neuron 64, 931–942, doi:S0896-6273(09)00886-1 [pii] 10.1016/j.neuron.2009.11.004 (2009).

60 Usher, M. & McClelland, J. L. The time course of perceptual choice: the leaky, competing accumulator model. Psychol Rev 108, 550–592 (2001).

61 Churchland, A. K. & Ditterich, J. New advances in understanding decisions among multiple alternatives. Current Opinion in Neurobiology 22, 920–926, doi:10.1016/j.conb.2012.04.009 (2012).

62 Bogacz, R. Optimal decision-making theories: linking neurobiology with behaviour. Trends Cogn Sci 11, 118–125, doi:10.1016/j.tics.2006.12.006 (2007).

63 Jiang, Z. D., King, A. J. & Moore, D. R. Topographic organization of projection from the parabigeminal nucleus to the superior colliculus in the ferret revealed with fluorescent latex microspheres. Brain Res 743, 217–232 (1996).

64 Graybiel, A. M. A satellite system of the superior colliculus: the parabigeminal nucleus and its projections to the superficial collicular layers. Brain Res 145, 365–374, doi:0006-8993(78)90870-3 [pii] (1978).

65 Asadollahi, A., Mysore, S. P. & Knudsen, E. I. Stimulus-driven competition in a cholinergic midbrain nucleus. Nat Neurosci 13, 889–895 (2010).

66 Knudsen. Control from below: the role of a midbrain network in spatial attention. Eur J Neurosci 33, 1961–1972 (2011).

67 Kothari, N. B., You, W.-K. & Mysore, S. P. Interactions between the superior colliculus and the lateral tegmental nucleus in the mouse. Society for Neuroscience (2018).

68 Deichler, A. et al. A specialized reciprocal connectivity suggests a link between the mechanisms by which the superior colliculus and parabigeminal nucleus produce defensive behaviors in rodents. Sci Rep 10, 16220, doi:10.1038/s41598-020-72848-0 (2020).

69 Ratcliff, R. & McKoon, G. The diffusion decision model: theory and data for two-choice decision tasks. Neural Comput 20, 873–922, doi:10.1162/neco.2008.12-06-420 (2008).

70 Roe, R. M., Busemeyer, J. R. & Townsend, J. T. Multialternative decision field theory: A dynamic connectionist model of decision making. Psychological Review 108, 370–392, doi:10.1037//0033-295x.108.2.370 (2001).

71 Deco, G., Rolls, E. T., Albantakis, L. & Romo, R. Brain mechanisms for perceptual and reward-related decision-making. Prog Neurobiol 103, 194–213, doi:10.1016/j.pneurobio.2012.01.010 (2013).

72 Louie, K., Grattan, L. E. & Glimcher, P. W. Reward value-based gain control: divisive normalization in parietal cortex. J Neurosci 31, 10627–10639, doi:10.1523/JNEUROSCI.1237-11.2011 (2011).

73 Furman, M. & Wang, X. J. Similarity effect and optimal control of multiple-choice decision making. Neuron 60, 1153–1168, doi:10.1016/j.neuron.2008.12.003 (2008).

74 Wong, K. F. & Wang, X. J. A recurrent network mechanism of time integration in perceptual decisions. J Neurosci 26, 1314–1328, doi:10.1523/JNEUROSCI.3733-05.2006 (2006).

75 Lo, C. C. & Wang, X. J. Cortico-basal ganglia circuit mechanism for a decision threshold in reaction time tasks. Nat Neurosci 9, 956–963, doi:10.1038/nn1722 (2006).

76 Roesch, M. R. & Olson, C. R. Neuronal activity in primate orbitofrontal cortex reflects the value of time. J Neurophysiol 94, 2457–2471, doi:10.1152/jn.00373.2005 (2005).

77 Kable, J. W. & Glimcher, P. W. The neural correlates of subjective value during intertemporal choice. Nat Neurosci 10, 1625–1633, doi:nn2007 [pii] 10.1038/nn2007 (2007).

78 Padoa-Schioppa, C. & Assad, J. A. Neurons in the orbitofrontal cortex encode economic value. Nature 441, 223–226, doi:10.1038/nature04676 (2006).

79 Machens, C. K., Romo, R. & Brody, C. D. Functional, But Not Anatomical, Separation of “What” and “When” in Prefrontal Cortex. Journal of Neuroscience 30, 350–360, doi:10.1523/Jneurosci.3276-09.2010 (2010).

80 Churchland, A. K. & Ditterich, J. New advances in understanding decisions among multiple alternatives. Curr Opin Neurobiol 22, 920–926, doi:10.1016/j.conb.2012.04.009 (2012).

81 Economo, M. N., Hansen, K. R. & Wachowiak, M. Control of Mitral/Tufted Cell Output by Selective Inhibition among Olfactory Bulb Glomeruli. Neuron 91, 397–411, doi:10.1016/j.neuron.2016.06.001 (2016).

82 Mori, K., Nagao, H. & Yoshihara, Y. The olfactory bulb: coding and processing of odor molecule information. Science 286, 711–715, doi:10.1126/science.286.5440.711 (1999).

83 Seger, C. A. & Miller, E. K. Category learning in the brain. Annu Rev Neurosci 33, 203–219, doi:10.1146/annurev.neuro.051508.135546 (2010).

84 Humphries, M. D., Stewart, R. D. & Gurney, K. N. A physiologically plausible model of action selection and oscillatory activity in the basal ganglia. J Neurosci 26, 12921–12942, doi:10.1523/JNEUROSCI.3486-06.2006 (2006).

85 Wang, L. & Krauzlis, R. J. Visual Selective Attention in Mice. Curr Biol 28, 676–685 e674, doi:10.1016/j.cub.2018.01.038 (2018).

86 Felsen, G. & Mainen, Z. F. Neural substrates of sensory-guided locomotor decisions in the rat superior colliculus. Neuron 60, 137–148 (2008).

87 Newsome, W. T., Britten, K. H. & Movshon, J. A. Neuronal correlates of a perceptual decision. Nature 341, 52–54, doi:10.1038/341052a0 (1989).

88 Chaure, F. J., Rey, H. G. & Quian Quiroga, R. A novel and fully automatic spike-sorting implementation with variable number of features. J Neurophysiol 120, 1859–1871, doi:10.1152/jn.00339.2018 (2018).

89 Rodriguez, A. & Laio, A. Machine learning. Clustering by fast search and find of density peaks. Science 344, 1492–1496, doi:10.1126/science.1242072 (2014).

90 Tibshirani, R., Walther, G. & Hastie, T. Estimating the number of clusters in a dataset via the gap statistic. J. R. Statist. Soc. B 63, 411–423 (2001).

